# Quantitative system drift

**DOI:** 10.1101/2025.09.17.676933

**Authors:** Carl Veller, Pavitra Muralidhar

## Abstract

We consider a biological system composed of multiple genetically variable components, the combined result of which is a quantitative trait under stabilizing selection for an optimal value. We show mathematically that, while the mean value of the system is ultimately constrained to remain near its optimum, the values of individual components are free to drift far from their initial values. Each component’s drift, though qualitatively similar to neutral drift, is slower by a factor that depends on the fraction of the system’s genetic variance contributed by the component. Our results provide a population-genetic basis for ‘system drift’, the concept that individual components of a biological system can evolve despite selective constraint on their combined product. A special case is a single polygenic trait under stabilizing selection, where our results predict that the mean genetic contributions to the trait of different subregions of the genome, such as the chromosomes, can drift despite constraint on the genome-wide genetic value. We explore the implications of this latter result for selection against interspecific hybrids and selection against turnovers of sex-determining systems. We further apply our general results to a continuous public goods game played between two species, where they predict that individual species’ contributions to a costly public good can drift freely. Finally, we show that symmetric mutation between alleles that increase and decrease components’ contributions to the system provides a weak long-term brake on components’ drift.

## 1 Introduction

In biological systems, multiple components combine to determine the outcome. The theory of system drift posits that the components of a system can undergo genetic evolution in spite of stabilizing selection to keep their combined product constant, because multiple configurations of the components can produce the same system outcome (True and Haag 2001). This theory has found widespread application, including in developmental genetics (True and Haag 2001; Sommer 2012; McColgan and DiFrisco 2024), gene-regulatory networks (Schiffman and Ralph 2022; Gaertner et al. 2023), and speciation (Barton 2001; Chevin et al. 2014; Schiffman and Ralph 2022; Schneemann et al. 2024).

Despite its importance as a general theory of the evolution of genotype–phenotype maps, system drift has received surprisingly little general population-genetic study. Population-genetic analyses of particular instances of system drift have been carried out. For example, Thompson et al. (2016) study how duplicate genes can drift in their individual expression levels despite stabilizing selection on their combined dosage, via successive substitutions of mutually compensatory mutations that alter each duplicate’s expression levels. In a similar successive-substitutions regime, a large literature has studied the consequences for hybrid fitness of independent substitutions that occurred in the parent populations and affect a trait under stabilizing selection (reviewed in Schneemann et al. 2024). Finally, Schiffman and Ralph (2022) study a model of a gene-regulatory network and show that its ‘wiring’ can evolve over time despite stabilizing selection on its product.

Here, we study a general model where the outcome of a system is a quantitative trait under stabilizing selection, and the system comprises some number of components that together determine its value. We show that the components will drift in their contributions to the system’s overall value, despite stabilizing on this value, as long as they present persistent genetic variation. This drift is qualitatively similar to neutral drift of the components’ contributions, as would occur if there were no selection on their combined product, but it is slower than neutral drift by a constant factor that we calculate. Our results thus provide a population-genetic basis for, and quantification of, the theory of system drift.

Because the theory of system drift has been invoked in a wide range of biological fields, to demonstrate the generality and versatility of our calculations, we apply them to three seemingly very different biological examples, one concerning reproductive isolation of species, one concerning transitions between sex-determining mechanisms, and one concerning cooperation in interspecies mutualisms.

## 2 Results

### 2.1 The basic model

There are *n* genetically variable components in a system, and their genetic bases are assumed not to overlap. These ‘components’ could be different species in a multispecies community, for example, or discrete components of a developmental system or molecular pathway, or even just distinct regions of the genome—such as the chromosomes—contributing to a quantitative trait like height. The components are assumed to combine additively to produce an overall quantitative trait that is under stabilizing selection. While our results do require that the overall trait, and thus individual components’ contributions to it, be quantitative, the assumption of additivity is not as restrictive as it might appear. If, for example, the components act in temporal sequence, it might be reasonable to assume that their effects combine multiplicatively, in which case rescaling the overall value of the system to be the logarithm of the original value rescues additivity of the components’ contributions. Additionally, while our model does require that multiple components of the system be genetically variable, it does not preclude genetically fixed components—the contributions of fixed components simply shift the optimal value that the genetically variable components must obtain. Therefore, without loss of generality, we ignore fixed components of the system in what follows. Finally, although we primarily focus on a model where the system has a single output under stabilizing selection, we note that biological systems often have many correlated outputs and can therefore be thought of more broadly as multivariate traits under multivariate stabilizing selection. In Appendix A2, we extend our model to consider systems with multiple outputs under stabilizing selection. But the principle of system drift and its connection to population-genetic models are easier to understand in the context of a system with a single output, and so we focus on this case in the Main Text.

Let 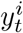 be the mean contribution of component *i* in generation *t*, and let 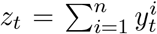 be the mean overall value of the system. We may arbitrarily set the optimal value of the system to be 0; the fitness of an individual with trait value *z* is then 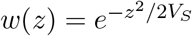, which can be approximated by 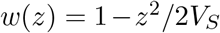 near *z* = 0. *V*_*S*_ calibrates the strength of selection on the value of the overall system; selection is stronger when *V*_*S*_ is smaller. Let *V*_*P*_ be the phenotypic variance of the system (typically ≪ *V*_*S*_), and let *V*_*G*_ be the genetic variance, with 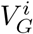 the contribution of component 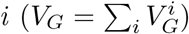. We assume these variances to be constant over the timescale we are interested in. The population comprises *N*_*e*_ diploid individuals.

### 2.2 Behavior of the mean value of the system

The mean value of the system *z*_*t*_ behaves as an Ornstein–Uhlenbeck (OU) process described by the stochastic differential equation (SDE)

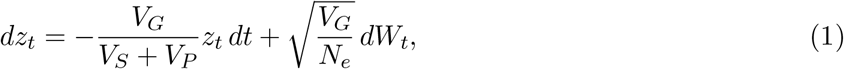

where *W*_*t*_ is a standard Brownian motion (e.g., Lande 1976, although in that paper, as is standard, the OU process for the mean value of a trait under stabilizing selection is cast in terms of an analogous partial differential equation (PDE) for the probability distribution over *z*_*t*_; it will turn out that, for our purposes, the SDE format is much more workable—see Discussion). The first term on the right-hand side of Eq. (1) is the deterministic change in *z*_*t*_ due to selection pulling it towards the optimum 0—it is negative if *z*_*t*_ *>* 0 and positive if *z*_*t*_ < 0. The system’s response to selection is faster if there is more genetic variance *V*_*G*_ for the value of the system and if the strength 1*/V*_*S*_ of stabilizing selection on that value is greater. The second term is a random perturbation to *z*_*t*_ due to genetic drift at polymorphic loci that affect the system’s value—it tends to be larger in magnitude if there is more genetic variance and if the population’s effective size is smaller.

From an initial value *z*_0_, the trajectory of *z*_*t*_ is described by

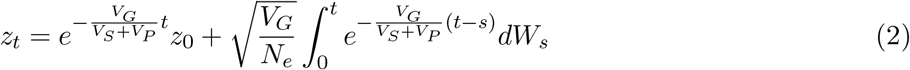

(e.g., Gard 1988, pg. 115). *z*_*t*_ thus approaches the optimum 0 exponentially (the first term in Eq. 2), but with random shocks due to genetic drift (the second term). The distribution of *z*_*t*_ across sample trajectories is normal:

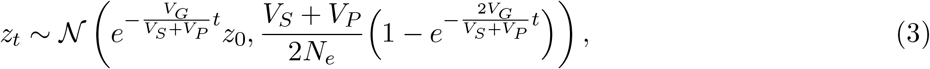

and converges on a timescale of ∼ (*V*_*S*_ + *V*_*P*_)*/V*_*G*_ generations to a long-term stationary distribution:

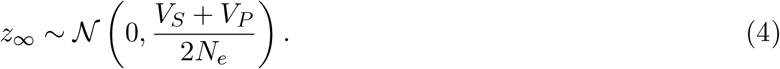

Thus, if the population size *N*_*e*_ is large, the mean value of the overall system is eventually contrained by selection to remain close to the optimum.

### 2.3 Behavior of the mean values of individual components of the system

The results above are standard to the literature on quantitative traits under stabilizing selection (e.g., Lande 1976; Simons et al. 2018; Hayward and Sella 2022). Our interest, however, is not in the behavior of the overall value of the system, but rather in the behavior of its constituent components. Are they constrained to remain in a narrow window around a particular value, or do they follow some other behavior?

We can decompose Eq. (1) into a system of coupled SDEs for the mean values of these components:

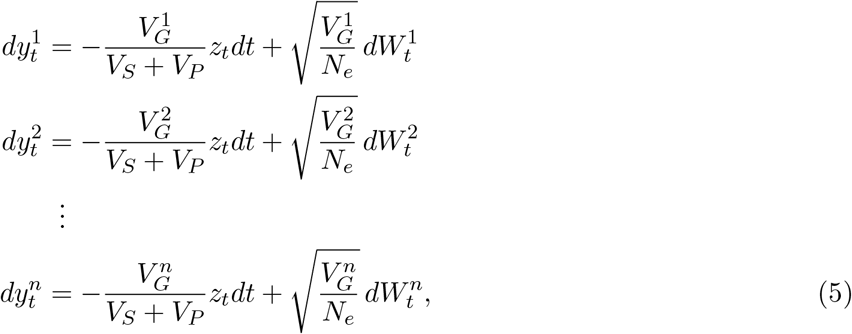

where the 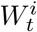 are independent standard Brownian motions. It is straightforward to verify that the equations in (5) sum to Eq. (1).^1^ The first terms on the right-hand sides of (5) indicate that every component is under selection to bring the mean value of the system towards the optimum 0; the common selection gradient, −*z*_*t*_*/*(*V*_*S*_ + *V*_*P*_), is larger in magnitude if the trait is far from the optimum and under strong stabilizing selection. The response of component *i* to this selection gradient is proportional to that component’s genetic variance 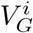. The second terms on the right-hand sides indicate that each component *i* is also subject to independent drift, which tends to cause deviations in the component’s mean value that are larger if the population size *N*_*e*_ is small and if the genetic variance for the component’s contribution to the system, 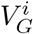, is large.

Our goal is to ‘decouple’ the equations in (5) to isolate the behavior of the mean contribution of each individual component *i* to the system. Let the fraction of the overall genetic variance of the system that is due to genetic variation in component *i* be 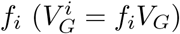, so that 1 − *f*_*i*_ is the fraction from all other components. Isolating the *i*-th equation in (5), and then summing all equations but the *i*-th (using the superscript −*i* for combined values of these other components), we find

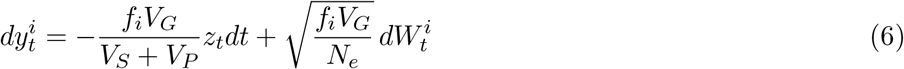

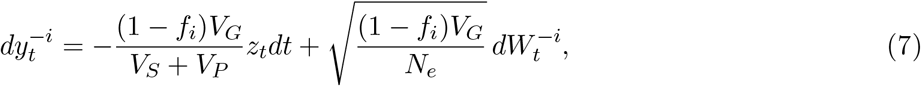

where 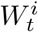 and 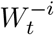 are independent standard Brownian motions. Dividing (6) by *f*_*i*_ and (7) by 1 − *f*_*i*_,

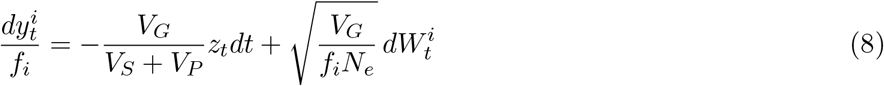

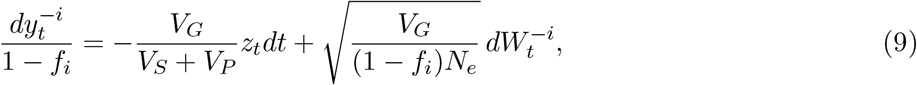

and subtracting (9) from (8),

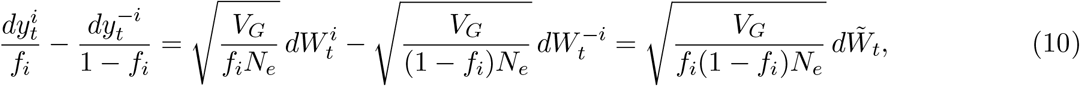

where 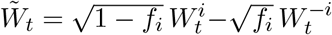 is a standard Brownian motion. Thus, the scaled difference between the mean contribution of component *i* and the mean contributions of all other components, 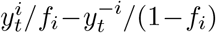, behaves as a Brownian motion. But 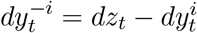, and so

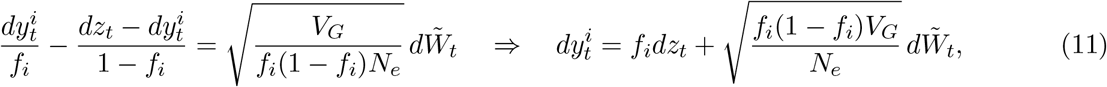

which we integrate:

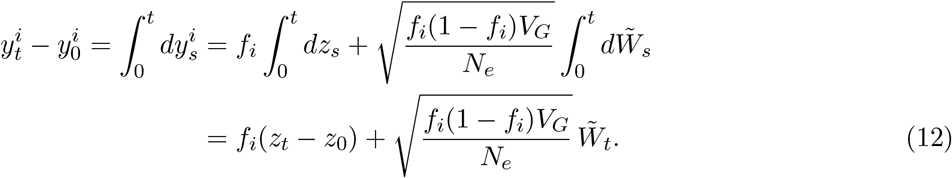

**Component drift is qualitatively neutral.** The trajectory of 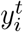, starting from its initial value 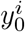, is therefore composed of two Gaussian processes. The first, *f*_*i*_(*z*_*t*_ − *z*_0_), is an OU process governing the convergence of the mean value of the overall system to its optimum. The second, 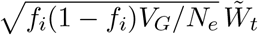 is a Brownian motion. In Appendix A1, we show that these two Gaussian processes are independent, so 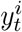 is itself a Gaussian process. Its mean in generation *t* is

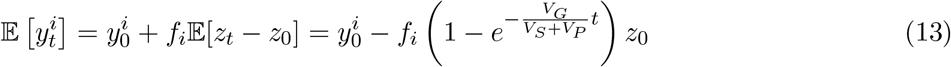

and its variance is

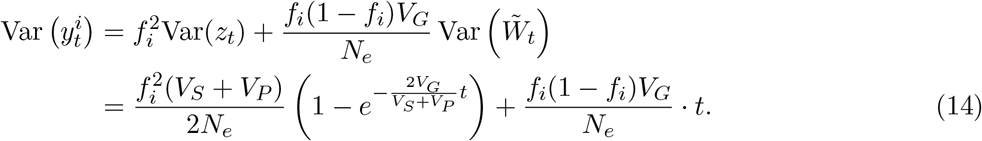

The mean and variance converge on a timescale of ∼ (*V*_*S*_ + *V*_*P*_)*/V*_*G*_ generations to 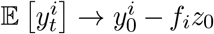 and

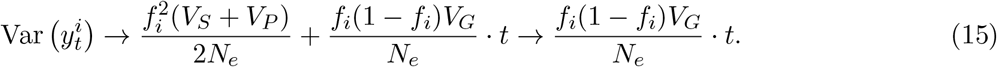

Therefore, the variance of 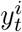 eventually grows linearly in time; that is, 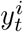 comes to behave as a Brownian motion. This occurs once the Brownian component of the variance in (15) dominates the OU component:

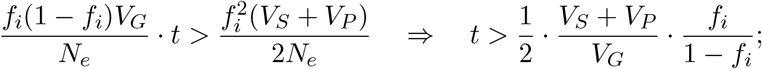

i.e., on the same timescale, ∼ (*V*_*S*_ +*V*_*P*_)*/V*_*G*_, as the convergence of the overall system’s OU process (Eq. 3) unless the component in question accounts for almost all of the genetic variance of the system (so that *f*_*i*_*/*(1 − *f*_*i*_) is very large).

In summary, despite the constraint that the mean value of the overall system must eventually remain close to its optimum, the mean values of the components of the system instead come to behave as Brownian motions, free to drift far from their initial values in a way that is qualitatively as if they were not under selection at all. Notice though that the rate of drift of 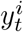, measured as its variance after *t* generations, would be *f*_*i*_*V*_*G*_*t/N*_*e*_ in the absence of selection. The Brownian rate in Eq. (14) is reduced from this neutral rate by a factor 1 − *f*_*i*_. This is the effect of selection on the overall system and, sensibly, if the component in question accounts for a large fraction of the genetic variance of the system (*f*_*i*_ ≈ 1), selection contrains this component to evolve more slowly.

The intuition for why the components drift continuously in their contributions to the system, despite selective constraint on the system’s value, lies in the persistent genetic variance that the components are assumed to show in our model. Consider a two-component system, and suppose that the mean contributions of the components to the system’s value are currently both zero, so that the system’s mean value is 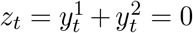, the optimum. Because the components show continuous genetic variation, some individuals in the population will have values for these components that deviate from zero but which nonetheless sum to zero—these individuals will be equally as fit as those at the population mean. The presence of such individuals distributed along the fitness ridge *y*^1^ + *y*^2^ = 0 guarantees drift along this ridge, i.e., correlated drift in the components’ mean values despite constancy of their sum. A similar intuition has been given by Schiffman and Ralph (2022).

For simplicity, in demonstrating the general possibility of system drift above, we have assumed that the output of the system is a single quantitative trait under selection. In reality though, systems—e.g., in development—will often have multiple, potentially correlated outputs. In Appendix A2, we generalize the calculations above to allow the components to contribute to multiple system outputs under selection. There, we show that our general result, that the components’ mean contributions to the system’s outputs come to behave as Brownian motions, is robust to this generalization.

### 2.4 Chromosome-scale drift under stabilizing selection

A particularly concrete case of the general model above is where the ‘system’ is any quantitative trait under stabilizing selection, and where the ‘components’ of the system are the genetic values contributed to the trait by nonoverlapping genomic subregions—such as the chromosomes—that together cover the entire genome. Note that the genomic subregions can constitute any partition of the genome and, in particular, need not be contiguous.

In this case, the calculations above imply that, while the genome-wide mean genetic value for the trait, *z*_*t*_, will ultimately remain near its optimum, the mean genetic values of different genomic subregions, 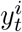, will be free to drift, with rates that depend on the fractions that they contribute to the total genetic variance for the trait (Fig. 1).^2^

**Figure 1:**
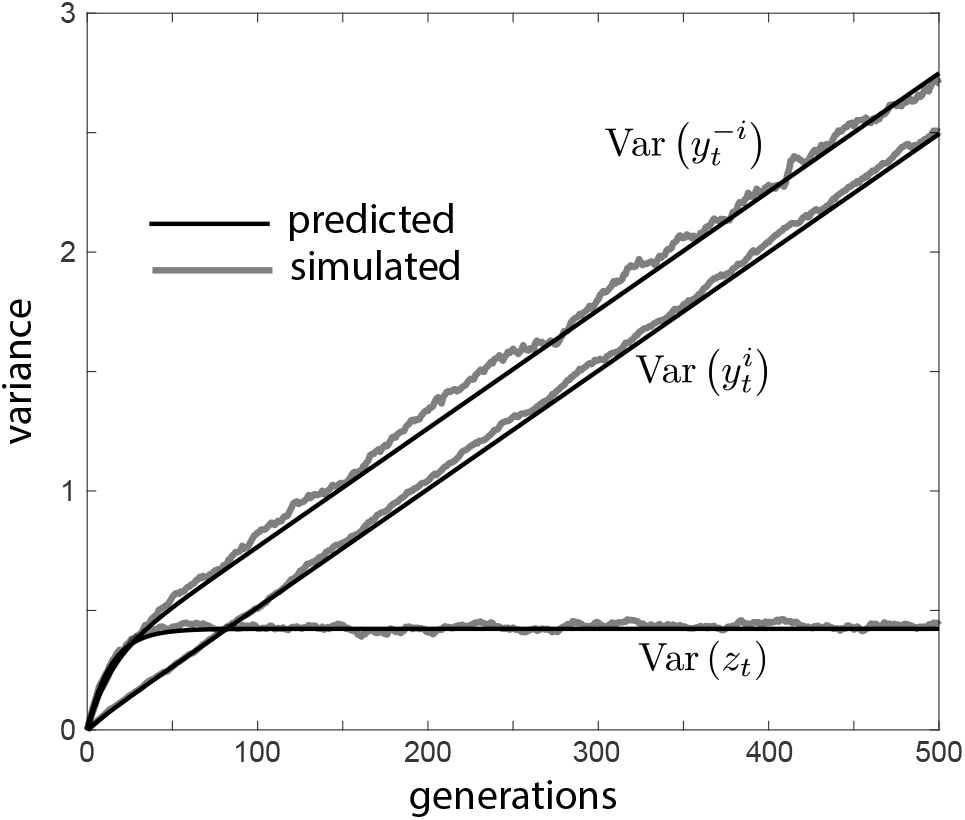
Chromosome-scale drift under stabilizing selection. Variance across evolutionary replicates of the mean genetic value of a genomic subregion *i*, its complement −*i*, and the entire genome for a quantitative trait under stabilizing selection. The population size is *N*_*e*_ = 10,000, and region *i* accounts for a fraction *f*_*i*_ = 0.2 of the genetic variance of the trait. Black lines show the predictions of Eq. (14) for regions *i* and −*i* and Eq. (3) for the whole genome. Grey lines show empirical variances across 2,000 replicate simulations in SLiM. In the SLiM simulations, there were 1,000 freely recombining loci, 200 in region *i* and 800 in region −*i*. A shared setup was generated by drawing minor-allele frequencies at these loci independently from a uniform distribution on [0.1, 0.3] and effects independently from a normal distribution with mean 0 and variance 1. The initial genetic variance *V*_*G*_ for this setup was calculated, and the strength of stabilizing selection was chosen so *V*_*S*_ = 25*V*_*G*_. This configuration was subjected to a 50-generation burn-in, the end-state of which was saved and used to initialize all 2,000 replicates.

This establishes a ‘mesoscale’ dynamics of the genetics of quantitative traits under stabilizing selection, occurring on a chromosome-level scale that is intermediate between the scale of the genome-wide dynamics (an OU process; Lande 1976 and above) and of the dynamics at individual loci (which resemble underdominant selection; Robertson 1956). These mesoscale dynamics, which we have shown to follow a Brownian motion (though see Section 2.8), are straightforwardly consistent with the genome-wide dynamics (since the system in Eq. 5, from which we have derived the mesoscale dynamics, sums to Eq. 1); their precise connection to the underdominant-like dynamics at individual loci is an avenue of future study.

A similar result has been obtained by Slatkin and Lande (1994) for the dynamics of the mean contribution of a single locus to a polygenic trait under stabilizing selection, in the case where mutation is stronger than selection at the locus so that it harbors many alleles (see their Eq. 4, and see also discussion in Lande 1975).

As noted above, in Appendix A2 we extend our general calculations to allow for multiple system outputs under stabilizing selection. Our results there imply, as a special case, that our finding that genomic subregions can drift in their contributions to a polygenic trait under stabilizing selection is robust to pleiotropic effects of the underlying genetic variants on other traits under stabilizing selection. This result will be relevant to our analysis of sex-chromosome turnover in Section 2.7.

### 2.5 Application 1: Reproductive isolation caused by stabilizing selection

The process of genomic subregion drift described above underlies the finding of Veller and Simons (2024) that, when two populations admix in unequal proportions, stabilizing selection on a quantitative trait generates selection against the minor-parent ancestry, even when the parent populations were (i) demographically identical, (ii) under the same regime of stabilizing selection, and (iii) recently diverged, such that they shared much of their genetic variation for the trait under selection (see also Slatkin and Lande 1994; Ragsdale 2025).

The logic is that, during their period of divergence, within each population A and B, the mean genetic value of a given genomic subregion happened to drift, with the complementary portion of the genome drifting in its value so as to keep the genome-wide value near the optimum. Because this subregion drift was independent in the two populations, a hybrid gamete with population-A ancestry in the one subregion and population-B ancestry in the complementary subregion will not necessarily have a mean genetic value close to the optimum (unlike gametes with the same ancestry in both portions). Hybrid gametes, and hence hybrid offspring, therefore show increased genetic variance and thus reduced fitness under stabilizing selection (Lande 1981). Integrating these reduced fitnesses across the distribution of hybrid ancestries in the admixed population in each generation after the admixture event, Veller and Simons (2024) find that stabilizing selection gradually purges the minor-parent ancestry from the admixed population over time. This argument of Veller and Simons (2024) rests on an assumption that the mean genetic value of a given genomic subregion can diverge substantially between the two populations despite both being under identical regimes of stabilizing selection. Our calculations above allow us to demonstrate and quantify this effect.

Suppose that, *t* generations ago, an ancestral population split into two descendent populations, A and B, of effective sizes 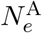 and 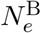. A quantitative trait has been under stabilizing selection of strength *V*_*S*_ in both populations, for a common optimal value that we arbitrarily set to 0. The genetic variance of the trait has been maintained at 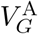 and 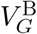 in the populations A and B respectively. The populations now mix in some proportions, and the admixed population remains under the same regime of stabilizing selection on the quantitative trait.

In the admixed population, a hybrid gamete is defined by a partition of its genome into two portions, the ancestry in one portion deriving from population A and the ancestry in the complementary portion deriving from population B. Consider one such partition of the genome into portions *i* and −*i*, which respectively account for fractions *f*_*i*_ and 1 − *f*_*i*_ of the genetic variance of a trait under stabilizing selection. We assume *f*_*i*_ to be the same for populations *A* and *B*, consistent, for example, with *f*_*i*_ being equal to region *i*’s fraction of total genome length. In the ancestral population, the mean genetic values of the two regions, 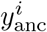 and 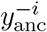, were co-adapted so that 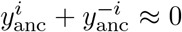. From Eq. (15), the mean genetic values of these regions in the two populations, after their period of divergence, are approximately

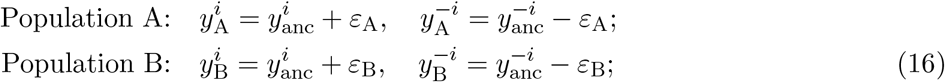

where *ε*_A_ and *ε*_B_ are independent and normally distributed:

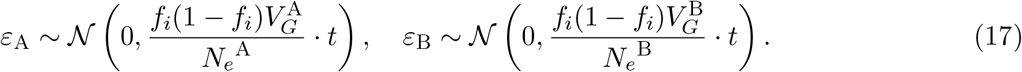

Let 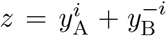 be the mean genetic value of a hybrid gamete in which the ancestry of genomic portions *i* and −*i* derives from populations A and B respectively. Since the fitness of an individual with trait value *x* is 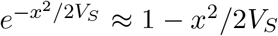, we are interested, for this particular partition of the genome, in the average value (across evolutionary replicates, or ‘plays of the tape’) of *z*^2^*/*2*V*_*S*_, the reduction in fitness due to the deviation of the genetic value of the hybrid gamete from its optimum:

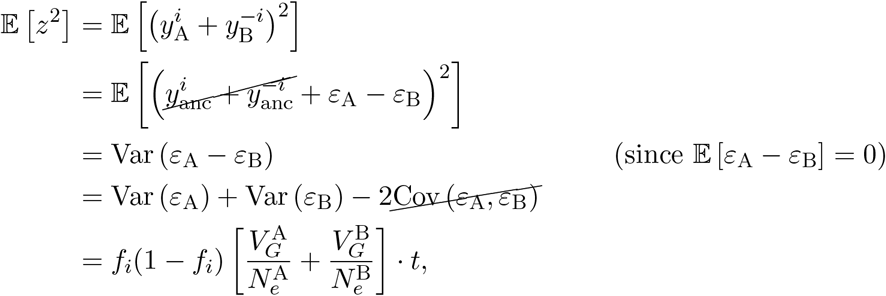

and so

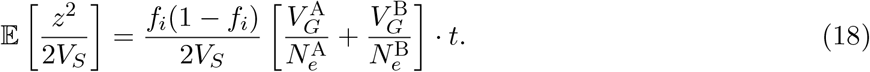

Note that a hybrid suffers reduced fitness equally, in expectation, whether the drift happened predominantly in parent population A or B, and, separately, whether it inherited predominantly A or B ancestry (by symmetry of the term *f*_*i*_(1 − *f*_*i*_)). Hybrids suffer greater fitness reductions, in expectation, if their ancestry fractions are closer to even, and given longer periods of divergence *t* of the parent populations.

In the case where the two populations are demographically identical, with common effective size *N*_*e*_ and genetic variance *V*_*G*_, Eq. (18) reduces to

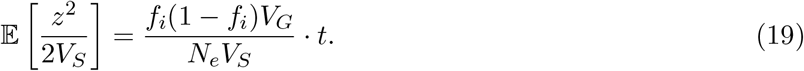

By a separate approach, Veller and Simons (2024) calculate that, if two demographically identical populations have diverged genetically at loci affecting a trait under stabilizing selection, then the expected fitness reduction from a gamete with proportions *f*_*i*_ of one ancestry and 1 − *f*_*i*_ of the other is

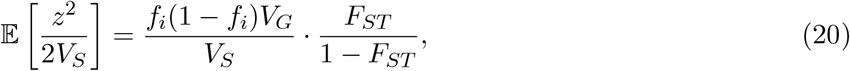

where *F*_*ST*_ is the degree of divergence at trait-affecting loci. Equating (19) and (20), we find that

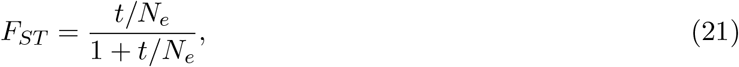

which gives, under stabilizing selection, the rate of accumulation of *F*_*ST*_ between the populations at trait-affecting loci. This rate is very similar to, though slightly lower than, the rate of accumulation of *F*_*ST*_ at neutral loci (∼ *t/N*_*e*_ on short timescales; 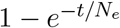 in general; e.g., Wakeley 2008).

Migration between the parent populations will naturally slow down the divergence of their mean genetic values in genomic portion *i*. In the case of symmetric sizes and genetic variances of the parent populations, we show in Appendix A3 that, if a fraction *m* of each parent population migrates to the other every generation during their period of divergence, then the difference between their mean genetic values in genomic portion 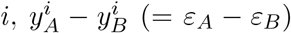, eventually comes to behave as an OU process with stationary distribution

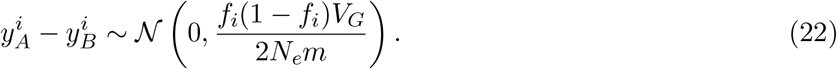

Under this stationary distribution, the reduced fitness of a hybrid that derives its ancestry in genomic portions *i* and −*i* from distinct parent populations is, in expectation,

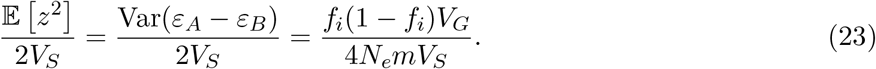

Thus, if migration is sufficiently common that 4*N*_*e*_*m* ≫ *V*_*G*_*/V*_*S*_, it will homogenize the parent populations sufficiently that hybrids between them will suffer negligible fitness reductions under stabilizing selection (see also discussion in Barton and Rouhani 1993).

### 2.6 Application 2: Public goods production in a mutualism

Species communities can also be thought of as systems, with the constituent species serving as interacting components of these systems. It is therefore interesting to consider whether such interactions can be subject to system drift. In this vein, suppose that two species interact in a mutualism, and that both require and are capable of producing some public good (an enzyme, say) which is released into their common environment and can be used by both species. For simplicity, we assume that the nature of the two species’ interaction is that they meet in pairs. For both individuals in an interacting pair, there is an optimal total amount *θ* of the good, and the strength of stabilizing selection *V*_*S*_ around this optimum is the same for the two species. However, the public good is also costly to produce, so that there is directional selection of intensity *S* for individuals to reduce their production.

We model this combination of selective forces by specifying that if, in a given interaction, the species-A interactant produces *Y* ^*A*^ of the good and the species-B interactant produces *Y* ^*B*^, the species-A individual has fitness 1 − *Y* ^*A*^*S* − (*Y* ^*A*^ + *Y* ^*B*^ − *θ*)^2^*/*2*V*_*S*_ (relative to other members of species A) while the species-B individual has fitness 1−*Y* ^*B*^*S*−(*Y* ^*A*^ + *Y* ^*B*^ − *θ*)^2^*/*2*V*_*S*_ (relative to other members of species B). Thus, if one individual among an interacting pair produces none of the good, its preference is for the other individual to produce *θ* of the good (in this sense, *θ* is the ‘optimal’ amount of the good—the amount that both individuals would like produced if they didn’t have to pay for it), while the other individual’s preference in this case is to produce less than *θ* (specifically, *θ* − *V*_*S*_*S*). In the language of Doebeli et al. (2004), this is a continuous nonlinear ‘snowdrift game’ with benefit function *B*(*x, y*) = 1 − (*x* + *y* − *θ*)^2^*/*2*V*_*S*_ and cost function *C*(*x*) = *Sx*, although in our case the game is played between two species rather than within one.

The two species have effective population sizes 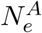 and 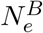, and phenotypic and genetic variances for production of the public good 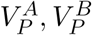 and 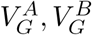 respectively. Let 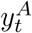 and 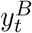 be the mean amounts produced by the two species, with 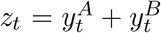 the overall mean amount. Then 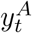 and 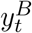 satisfy the coupled SDEs

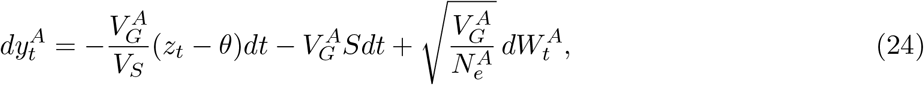

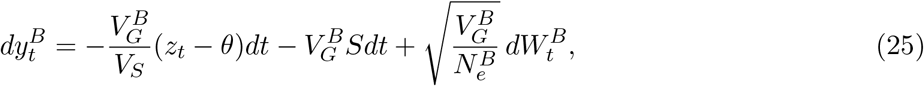

where 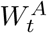 and 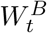 are independent standard Brownian motions. We have assumed that 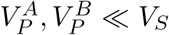 in simplifying the coefficients 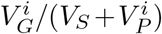 to 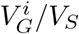 in Eqs. (24) and (25); the full selection gradients are derived in Appendix A4. If there are more than two species in the mutualism, Eq. (25) can be interpreted as governing the sum of the contributions of all species other than A, in a manner similar to Eq. (7).

Summing Eqs. (24) and (25), we find that the mean overall production of the public good, 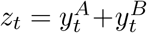 obeys

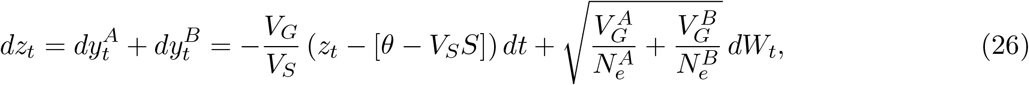

where 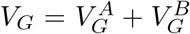 and *W*_*t*_ is a standard Brownian motion. Eq. (26) describes an OU process, with the mean overall production *z*_*t*_ evolving towards the v alue *z*^*∗*^ = *θ* − *V*_*S*_*S*. The stationary distribution of *z*_*t*_ is normal with mean *z*^*∗*^ and variance 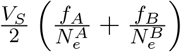, where 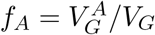 and 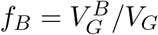.

To determine the behavior of the mean contribution of each species in the mutualism, we can use methods similar to those in Section 2.3 to scale and subtract Eqs. (24) and (25), finding

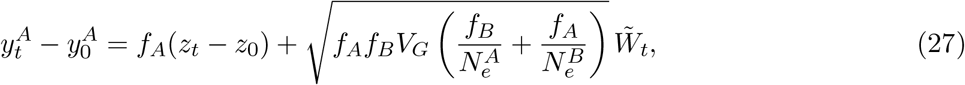

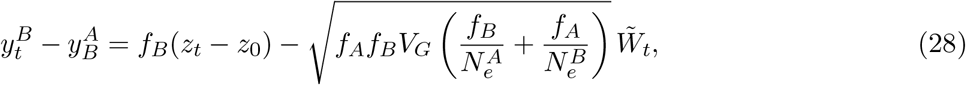

where 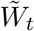 is a standard Brownian motion.

The two terms on each of the right-hand sides of Eqs. (27) and (28) are Gaussian processes and (following the same logic as in Appendix A1) they are independent; the mean contributions of the two species at time *t*, 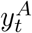 and 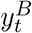, are therefore normally distributed, and their variances that tend to

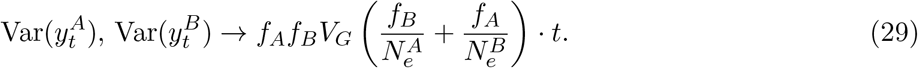

That is, the two species’ mean contributions to the public good come to behave as Brownian motions, despite being under directional and stabilizing selection.

How do the intrinsic rates of drift within the two populations affect their respective outcomes in the mutualism? To isolate the role of population size, assume 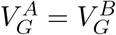 (i.e., *f*_*A*_ = *f*_*B*_ = 1*/*2). Then

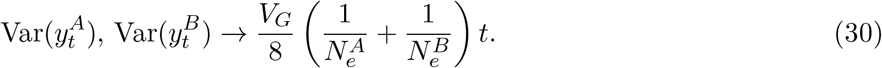

That is, the rate of drift of the two species’ mean contributions is governed by the harmonic mean of their population sizes, and is therefore predominantly determined by the smaller population. An appealing metaphor is that the smaller population is on a drunkard’s walk, and the larger population, sober though tethered to the drunkard, must counterstep to keep the pair on target; in doing so, however, the sober member of the pair also looks drunk.

A counterintuitive implication of the fact that the two species’ mean productions of the public good drift randomly is that, owing to the fitness cost of producing the good, their mean fitnesses also drift randomly. We anticipate that this result will be sensitive to certain symmetries that we have assumed in our model, such as the equal strengths of selection (both stabilizing and directional) in the two species, their agreement on the optimal production *θ* of the public good, and the linearity of directional selection against production. We will explore the effects of relaxing these assumptions elsewhere.

### 2.7 Application 3: Stabilizing selection as a barrier to sex chromosome turnover

Muralidhar (2024) has shown by whole-genome simulations that stabilizing selection on polygenic traits, encoded across sex chromosomes and autosomes, can act as a barrier to transitions between sex chromosome systems. The reason is that sex chromosome turnovers necessarily create novel sexual genotypes (such as YY males or XY females), and these novel sexual genotypes disrupt the genome-wide coadaptation of the chromosomes in setting traits under stabilizing selection to their optima. The theory requires that allelic effects in males and females be imperfectly correlated, and that recombination between the X and Y (or Z and W) chromosomes be shut down along a portion of their length. Since, in this theory, alleles have pleiotropic effects on the male and female trait value, our extension of the general calculations above to allow for pleiotropic effects of alleles on multiple traits under stabilizing selection (Appendix A2) allows us to provide some quantification of the theory.

We consider a randomly mating male-heterogametic (XX/XY) species of effective size *N*_*e*_. A quantitative trait is under stabilizing selection of strength 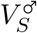 in males and 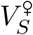 in females. We mark the male and female trait optima as zero.^3^ Let 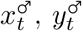, and 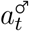 be the mean haploid genetic values of the X, the Y, and the autosomes for the trait in males in generation *t*, and let 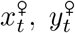, and 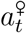 be the analogous values for the trait in females (note that, even though no females carry a Y chromosome in the initial male-heterogametic system, the Y chromosome nonetheless harbors a latent effect on the female trait, which would be expressed if some females came to carry Y chromosomes). Ignoring linkage disequilibrium, in the initial heterogametic system, the mean trait values in males and females in generation *t* are 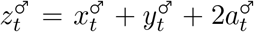 and 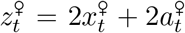. Define the column vectors 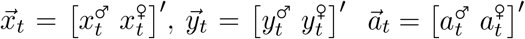, and 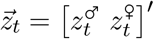.

The associated haploid genetic variances are 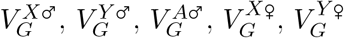, and 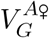, and the haploid genetic covariances between the male and female trait are 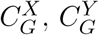, and 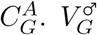 and 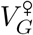 are the overall diploid genetic variances in males and females, and *C*_*G*_ is the overall diploid genetic covariance between the male and female trait values. We collect these genetic variances and covariances in the matrices

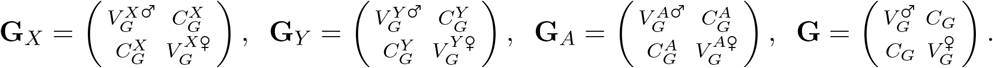

In the initial male-heterogametic system, 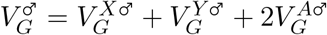 and 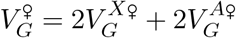.

We assume that the X and Y do not recombine in males; if recombination is shut down along only a portion of the X and Y, then their recombining portion can be considered autosomal. We further assume that the Y chromosome is nondegenerate, which in our case means that the genetic variance it contributes to the trait is comparable to that contributed by the X chromosome and there is no dosage compensation for the X chromosome in males (it contributes its haploid value).

In the initial male-heterogametic system, the mean contributions of the various regions of the genome to the male and female trait value behave according to the system of SDEs:

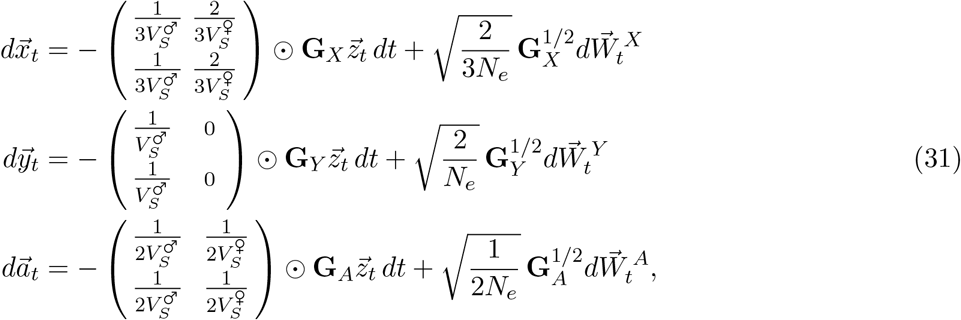

where ⊙ is element-wise multiplication and 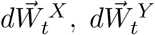, and 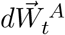 are independent two-dimensional standard Brownian motions. The selection matrices on the right-hand sides in Eq. (31) are derived in Appendix A5; the fractions in them reflect the fractions of time spent by the various genomic regions in males (left columns) versus females (right columns). Note that the nature of the genetic variances and covariances of the Y are qualitatively different to those of the other genomic regions: while the other regions comprise a large number of recombining loci, with genetic variances and covariances summed across these loci, the Y chromosome can be thought of as a single nonrecombining locus, the genetic variance at which is furnished by a large number of alleles. Despite this difference, the Y chromosome’s genetic variances and covariances enter Eq. (31) in a qualitatively similar way to those of the other regions.

Owing to the parametric complexity of (31) and, in particular, the non-proportionality of the matrix coefficients in the selection terms, we will not attempt to decouple the system analytically as we did earlier for simpler systems. However, we can iterate the system numerically to observe the behavior of the mean contributions of the various regions of the genome to the male and female trait values.

Fig. 2 shows trajectories of the variances of these contributions over time, averaged across replicate whole-genome simulations in SLiM and across replicate simulations of the SDEs in Eq. (31). The simulations in SLiM are seeded with particular initial genetic variances and intersex covariances for the X, Y, and autosomes. The genetic variances and covariances do not change much in these simulations over the relatively short timescale shown in Fig. 2, allowing us to predict the trajectories of the variances of the mean contributions of these regions over this timescale by substituting the initial genetic variances and covariances into Eq. (31). The contributions of the various regions of the genome are seen to drift over time, despite constraint on their combined value in both males and females, and this drift is well predicted by Eq. (31) on the timescale shown in Fig. 2. Moreover, the growth of the variances of these regions’ contributions tends to linear, consistent with these contributions behaving as Brownian motions, as for the simpler case where all genomic regions are autosomal, studied above.

**Figure 2:**
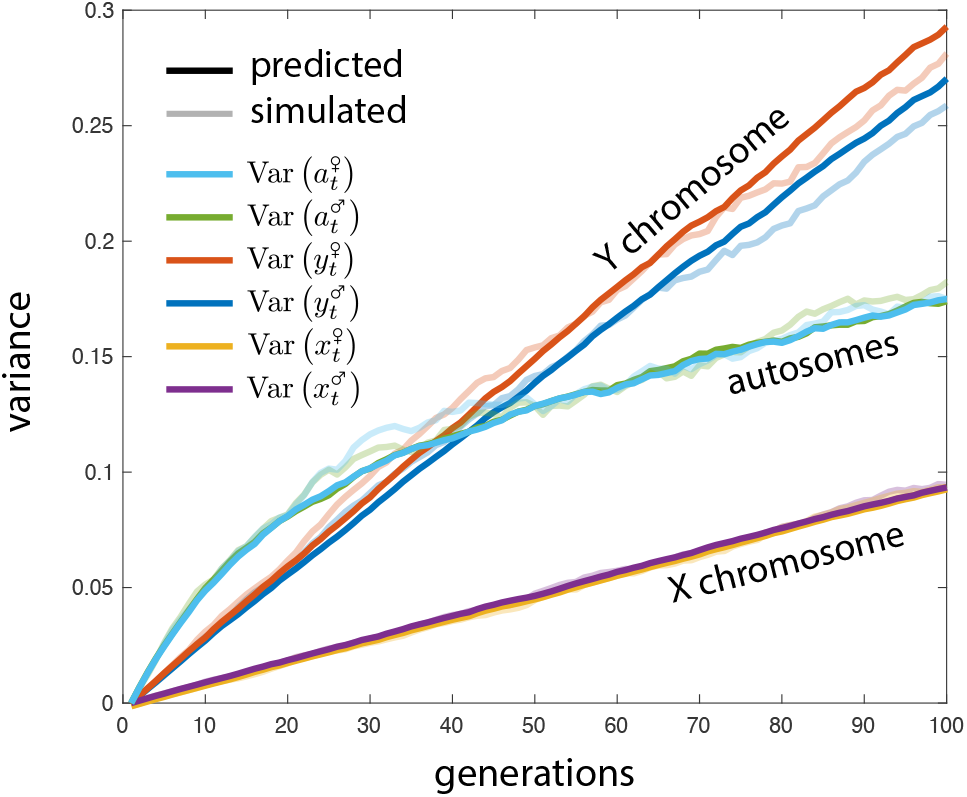
Sex chromosome drift under stabilizing selection. Variance across evolutionary replicates of the mean haploid genetic values of the X chromosome, the Y chromosome, and the autosomes for the male and female values of a trait under stabilizing selection. Solid lines are predictions based on Eq. (31). Faint lines are empirical variances across 1,000 replicate simulations in SLiM. In these simulations, *N*_*e*_ = 10,000, and there are initially 1,000 polymorphic loci, 100 on the sex chromosomes (both X and Y) and 900 on the autosomes. Loci recombine freely, except between the X and Y. A shared simulation setup was generated by drawing minor-allele frequencies independently from a uniform distribution on [0.1, 0.3] and the effects of the minor alleles on the male and female trait independently from a bivariate normal distribution with variance 1 and correlation 0.9. A resampling procedure was used to ensure that 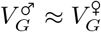 and that the regional G matrices scaled with genomic length (**G**_*X*_ **≈ G**_*Y*_ **≈ G**_*A*_*/*9). The strength of stabilizing selection in sex *k* was 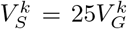. The chosen configuration was subjected to a 50-generation burn-in and the end state saved; all replicates were initialized from this saved state.

On longer timescales, such as those simulated by Muralidhar (2024), the genetic variances and covariances of the various regions will settle to equilibrium values determined by a balance between mutation, selection, and drift. On such timescales, the rate of drift of the contributions of these regions to the trait under selection will depend on the mechanisms maintaining their genetic variance, and in particular will depend on the nature of the mutations that replenish this variation as it is depleted by selection.

### 2.8 Mutation and the long-term evolution of a system’s components

The results above suggest that, although the mean value of a system under stabilizing selection is ultimately constrained to remain near its optimum, the mean contributions of individual components come to behave as Brownian motions, and are therefore free to drift far from their initial values. Here, we explore a long-term brake on this process: mutation. The idea is that components that have drifted to very large positive contributions to the trait have done so by the increase in frequency, and in some cases fixation, of positive-effect alleles at loci that affect the component’s contribution to the system. If mutation rates are symmetric between positive and negative alleles at component-affecting loci (including loci currently fixed for one or the other allele), the excess frequencies of positive alleles for this component present excess opportunities for mutation to negative alleles, and so mutation will tend to decrease the component’s mean contribution to the system. Similarly, components that have drifted to negative contributions to the system’s value will present excess opportunity for positive mutations, so mutation will tend to increase their mean contributions.

In Appendix A6, we introduce into our general calculations a symmetric per-generation per-locus mutation rate *μ* between positive- and negative-effect alleles at component-affecting loci. Assuming *μ* to be small, we show that the system’s mean value *z*_*t*_ still behaves as an OU process with long-term mean 0 and variance approximately

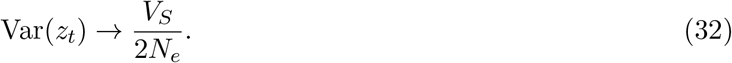

However, each component *i*’s mean contribution to the system, 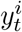, now comes to behave not as a Brownian motion, as in the case without mutation, but instead as an OU process with long-term mean 0 and variance approximately

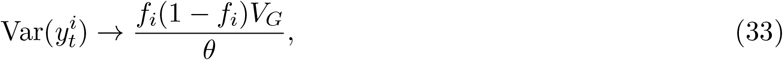

where *θ* = 4*N*_*e*_*μ* is the population-scaled per-locus mutation rate and *f*_*i*_ is the fraction of the system’s genetic variance contributed by component *i*.

The variance of 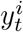 is thus bounded by mutation in the long run. While this implies a degree of constraint on the evolution of the mean values of the individual components of the system, this constraint is likely to be weak relative to the constraint on the mean genetic value of the overall system, and so there will still be substantial scope for system drift. Under stabilizing selection, in mutation–selection–drift balance in the large-population limit, *V*_*G*_ ≈ 4*V*_*S*_*Lμ*, where *L* is the number of loci at which mutations affect any of the components in the system (e.g., Turelli and Barton 1990). Substituting this value into Eq. (33), we find that the equilibrium variance of component *i*’s mean contribution to the system is *f*_*i*_(1−*f*_*i*_)*LV*_*S*_*/N*_*e*_. Compared with the variance of the overall system’s mean value, *V*_*S*_*/*2*N*_*e*_, the component-specific variance is larger by a factor 2*f*_*i*_(1 − *f*_*i*_)*L*. Since *L* ≫ 1 for a polygenic system, this factor will be very large unless component *i* contributes almost none (*f*_*i*_ ≈ 1*/L*) or almost all (*f*_*i*_ ≈ 1 − 1*/L*) of the system’s genetic variance.

For a concrete example, consider the contribution of an arbitrarily chosen half of the genome (*f*_*i*_ = 1*/*2) to a polygenic trait under stabilizing selection. This case is relevant, for example, to the fitness of second-generation hybrids in Application 1 above. In this case, the equilibrium variance of half the genome’s mean genetic value is larger than the equilibrium variance of the genome-wide mean by a factor *L/*2. For a highly polygenic trait, this factor will be extremely large (e.g., *>* 1,000 for human height). The mean genetic value of half the genome thus drifts across a very wide range of values over time, relative to the narrow range of values to which selection confines the genome-wide mean.

## 3 Discussion

We have shown that, when multiple components contribute to a system, the output of which is a quantitative trait under stabilizing selection, the components’ contributions to the system’s value will drift over time as long as they present persistent genetic variance. Each component drifts qualitatively as if the system is not under selection at all; quantitatively, however, its rate of drift is slower than neutral by a factor equal to one minus the fraction of the system’s genetic variance contributed by the component in question. Our results provide a population-genetic basis for, and quantification of, the theory of system drift (True and Haag 2001).

Our conclusions depend crucially on the assumption that the components show persistent genetic variance over time. This assumption is consistent with a modern understanding of the genetic architecture of quantitative traits, which association studies and other methods have shown usually to be affected by mutations at very many loci, presenting a large mutational target that ensures perpetual replenishment of genetic variation despite its depletion by selection (Sella and Barton 2019).

The theory of system drift was first proposed in the context of developmental biology (True and Haag 2001), and discussions of the theory in this context often seek to distinguish it from the concept of neutral genetic drift (e.g., True and Haag 2001; McColgan and DiFrisco 2024). Our results show that this conceptual distinction is unnecessary: as long as the system shows persistent genetic variance, system drift will occur via genetic processes that are qualitatively the same as neutral drift.

To make this connection clear, consider a particularly well-studied case of developmental system drift: the regulatory elements underlying expression of the *even-skipped* gene in *Drosophila*, a key factor controlling segmentation in early embryonic development (True and Haag 2001). Expression of *even-skipped* in the “stripe 2” region of the embryo is controlled by a short enhancer that integrates inputs from multiple transcription factors. Although the spatial pattern of stripe 2 expression is highly conserved across *Drosophila* species, comparative work (Ludwig et al. 1998, 2000) has revealed that the number, arrangement, and identity of transcription factor binding sites within the enhancer differ substantially across species. Under a traditional conception of developmental system drift, this divergence might be attributed to successive, discrete genetic substitutions at the enhancer in different lineages. However, evidence for intraspecific genetic variation in these regions (Ludwig and Kreitman 1995; Ludwig et al. 2000) suggests that the underlying regulatory components of this system might also have evolved through the continuous drift process described in this work.

### 3.1 Chromosome-scale dynamics of complex traits under stabilizing selection

One application of our general results to which we have paid special attention is where the system is a complex trait under stabilizing selection and the components of the system are the contributions to this trait of different regions of the genome, such as the chromosomes. In this case, our results imply that the mean genetic values of the different genomic subregions are free to drift, despite the overall trait value remaining close to its optimum. This reveals a ‘mesoscale’ genetic dynamics of complex traits under stabilizing selection, intermediate to the genetic dynamics at individual loci (which resemble weak fitness underdominance; Robertson 1956) and the dynamics of the genome-wide mean genetic value (an OU process; Lande 1976). The rate of a given subregion’s drift is slower than the neutral rate, by a factor equal to one minus the fraction of the trait’s genetic variance contributed by the subregion in question (which in practice will often be well proxied by the subregion’s fraction of total genome length; Yang et al. 2011). We have demonstrated two evolutionary consequences of these mesoscale dynamics, one for reproductive isolation of divergent populations and one for sex chromosome turnover, and we anticipate that the mesoscale dynamics will be found to have many other evolutionary consequences.

Importantly, and consistent with our discussion of the regulation of *even-skipped* above, the mesoscale dynamics do not preclude the fixation of alleles at individual loci. As long as there is persistent genetic variance for a genomic subregion’s contribution to the trait under selection, its mean contribution to the trait will drift. This will be true at any timescale, and in particular it will be true throughout the course of turnover of the genetic basis of variation in the trait, as ancestrally polymorphic loci become fixed and new polymorphisms are generated at other loci by mutation.

This property distinguishes our analysis from previous studies of long-term system drift in the mutation-limited regime. In this regime, trait-affecting mutations are assumed to be very rare, such that the evolution of a trait under stabilizing selection proceeds via successive substitutions of trait-affecting mutations in an otherwise monomorphic population. Unless the population starts far from its optimal trait value, some of these substitutions must occur against the grain of selection—indeed, once the population is monomorphic for an optimal genotype, any trait-affecting mutation is deleterious. That some substitutions must be deleterious constrains the rate and scope of system drift in the mutation-limited regime: either these mutations must have very small effects on the trait or the population must be very small (so that the mutations are nearly neutral). In contrast, our analysis shows that if the population is not monomorphic for the trait under stabilizing selection, and instead shows persistent genetic variance, then system drift occurs as an inevitable and qualitatively neutral process that furthermore permits, in its course, substitution of trait-affecting alleles.

One context to which system drift has often been applied is hybrid fitness and speciation (e.g., Barton 2001; Chevin et al. 2014; Schneemann et al. 2024). The scenario typically studied is where two populations split from an ancestral population and, under stabilizing selection on one or more quantitative traits, diverge at trait-affecting loci. When the populations come into secondary contact and hybridize, some hybrids receive unbalanced combinations of trait-affecting alleles from the two populations and suffer reduced fitness. Veller and Simons (2024) show that the mean fitness reduction to hybrids of any ancestry composition depends on the degree of divergence between the populations at trait-affecting loci, as measured by *F*_*ST*_ at these loci. In the mutation-limited regime, this divergence mjst occur via independent substitutions at trait-affecting loci in otherwise monomorphic populations and, unless the ancestral population was far from the trait optimum, some of these substitutions must be against selection (Chevin et al. 2014). This requirement limits the rate at which *F*_*ST*_ can accumulate between the populations in the mutation-limited regime, and thus limits the rate at which reproductive isolation can build up between them. In contrast, when the populations show persistent genetic variation for the traits under selection, our calculations here reveal that *F*_*ST*_ at trait-affecting loci accumulates at close to the neutral rate (Eq. 21). Our theory of subregion drift therefore entails more rapid accumulation of reproductive isolation between divergent populations.

Furthermore, because subregion drift permits fixation of alleles at trait-affecting loci, our theory covers both short-term and long-term divergence of the parent populations. It can therefore predict the average fitness of hybrids between recently diverged populations that still share much of their genetic variation for traits under stabilizing selection, as well as of hybrids between long-diverged populations that have evolved separate genetic bases of the traits under stabilizing selection. In this sense, our analysis here and in Veller and Simons (2024) of hybrid fitness under stabilizing selection and the subsequent purging of introgressed DNA in admixed populations is both qualitatively different from, and more general than, previous analyses restricted to the mutation-limited regime.

The assumption that distinguishes our analysis is that there is persistent genetic variation for traits under stabilizing selection. As we have emphasized, this is the expectation for traits with highly polygenic genetic architectures (Lande 1975), which genome-wide association studies and other methods are revealing to be the norm (Sella and Barton 2019).

### 3.2 Stochastic versus partial differential equations in population genetic modeling

In deriving our results, we have modeled the diffusion of components’ and the system’s mean genetic values using stochastic differential equations (SDEs; e.g., Eqs. 1 and 5). It is standard in population genetics to model diffusions, both of allele frequencies and of mean genetic values, instead in terms of partial differential equations (PDEs) governing probability distributions over their values (e.g., Lande 1976; Ewens 2004; though see Balick 2023, Bertram and Shafiei 2025, and Cope et al. 2025 for recent exceptions). While the SDE and PDE approaches are equivalent given appropriate boundary conditions, and while it is therefore in principle an arbitrary choice which to use, in practice we have found SDEs to be more workable for our purposes.

Consider, for example, the relationship between the equations in (5), which describe the change in the components’ mean values over time, and Eq. (1), which describes the change in the overall system’s mean value over time. To obtain the SDE for the system’s mean value (1) from the SDEs for the components’ mean values (5), we simply sum them. We could alternatively have written these equations in terms of their PDE analogs. But then to obtain the PDE for the overall system’s mean value from the PDEs for the components’ mean values would require computing successive convolutions of the probability distributions of the components’ values, a substantially less intuitive and more tedious task than simply suming a set of SDEs. We therefore propose that, for certain applications of diffusion theory in population genetics, SDE representations might be more practical (and, for pedagogical purposes, perhaps more intuitive) than PDE representations.

## Acknowledgements

We are grateful to Daniel Cooney, Graham Coop, Serena Debesai, Martin Nowak, Yuval Simons, Matthias Steinrücken, and Joe Thornton for helpful discussions, and to Graham Coop for helpful comments on the manuscript.

## A1 The OU and Brownian processes in Eq. (12) are independent

Eq. (12) in the Main Text describes the trajectory of the mean contribution of component *i* of the system, 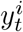, starting from its initial value 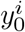:

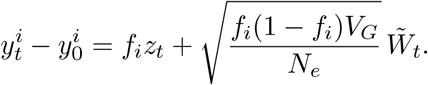

We want to show that the two processes on the right-hand side are independent, which follows if *z*_*t*_ and 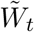 are independent. Defining 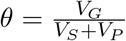 for brevity, recall from Eq. (2) that

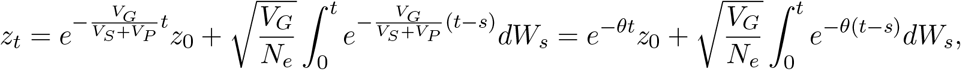

so in fact we need only show that 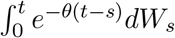 and 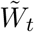 are independent. To do so, we will first show that 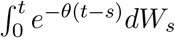 and 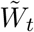 are jointly bivariate Gaussian, and then show that they are uncorrelated.

Since Eqs. (6) and (7) must sum to Eq. (1),

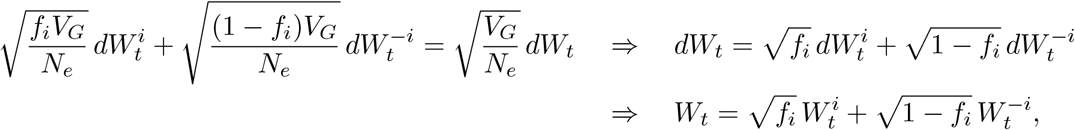

and from Eq. (10),

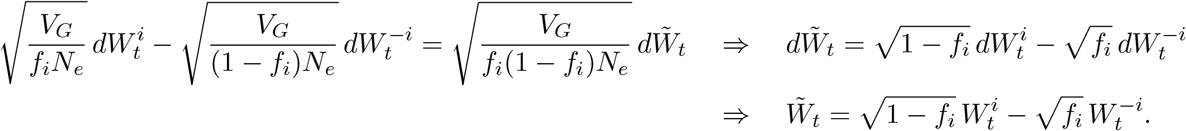

To show that 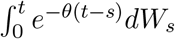 and 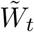 are jointly bivariate Gaussian, it is sufficient to show that any linear combination of them is Gaussian. For any constant factors *A* and *B*,

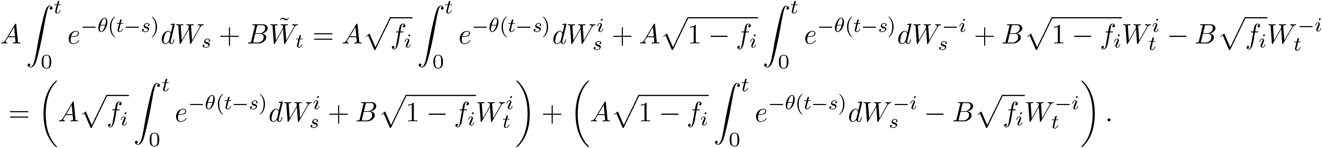

The two bracketed terms in the last line are independent Gaussian processes, since they are linear functionals of the independent Brownian processes 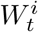 and 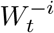 respectively. Their sum is therefore Gaussian.

We now show that the covariance of 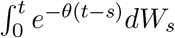 and 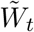 is zero:

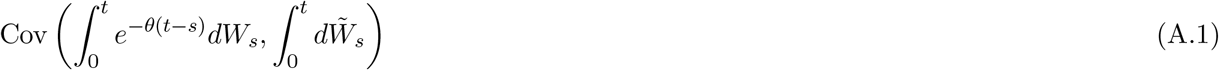

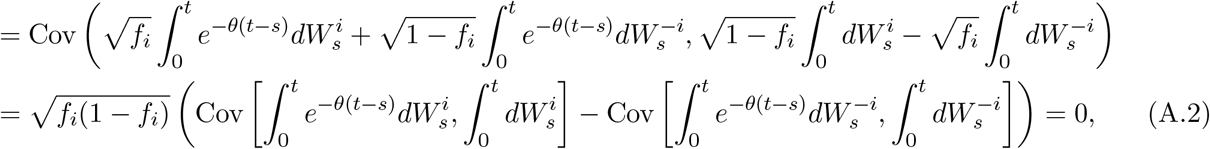

where we have used the fact that 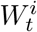 and 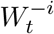 are independent. The covariances in Eq. (A.2) are straightforward to resolve using Itô isometry, but the final equality follows more simply from the symmetry of 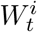 and 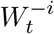.

Since 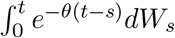 and 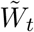 are bivariate Gaussian with zero covariance, they are independent.

## A2 Pleiotropy

The results in the Main Text assume that the system’s output is a single quantitative trait under stabilizing selection. Here, we extend our calculations in the Main Text to allow for systems with multiple, potentially correlated outputs under stabilizing selection, with alleles that affect the components on the system therefore having pleiotropic effects on these outputs. We frame our results in terms of multiple quantitative traits under stabilizing selection, focusing attention on the behavior of the mean contributions of different subregions of the genome to each trait, as in Main Text Section 2.4; however, our results should be interpreted broadly as applying to systems with multiple outputs under stabilizing selection.

We assume that there are *m* traits under stabilizing selection, with 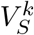 the strength of selection on trait *k*, and 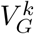 and 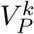 the overall genetic and phenotypic variance for trait *k* (note that the use of the superscript 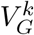 here, for the genetic variance of trait *k*, is distinct from the usage in Main Text Section 2.3, where it denoted the contribution of component *k* to the system’s genetic variance). We may arbitrarily set the optimal value of each trait to be 0. We focus on the contributions of *n* ‘components’ to these traits, with *y*^*ik*^ the mean contribution of component *i* to trait *k* at time *t*. The components are assumed to be genetically distinct, in that their genetic variation is underlain by distinct sets of loci in the genome. We denote by 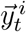 the *m ×* 1 vector of the mean contributions of component *i* to the *m* traits, with 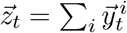 the resulting vector of mean overall genetic values of the *m* traits.

The genetic bases of the traits overlap, and alleles’ effects on the multiple traits are correlated. We assume the effects of each allele on the various traits are chosen from some multivariate distribution, independently at each of many loci, with correlation *ρ*_*jk*_ between traits *j* and *k*. Note that this is the correlation observed after many generations of selection, and is not the same as the correlations observed for new mutations since, for example, selection will preferentially cull alleles that have large effects on multiple traits (Simons et al. 2018). We further assume that each component is sufficiently polygenic that its empirical correlation of allelic effects on each pair of traits *j* and *k* is very close to *ρ*_*jk*_.

Under these assumptions, component *i* accounts for the same fraction, which we denote *f*_*i*_, of the genetic variance 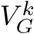 of every trait *k*, and this is also the fraction contributed by component *i* to the genetic covariance 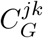 between every pair of traits *j* and *k*. Denote by **G** the *m × m* genetic variance–covariance matrix, of which the *k*-th diagonal element is 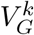 and the *j, k*-th off-diagonal element is 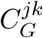. We assume **G** to be positive definite and approximately constant over the timescale we are interested in. Over this timescale, the mean contributions of the components to the traits then evolve according to

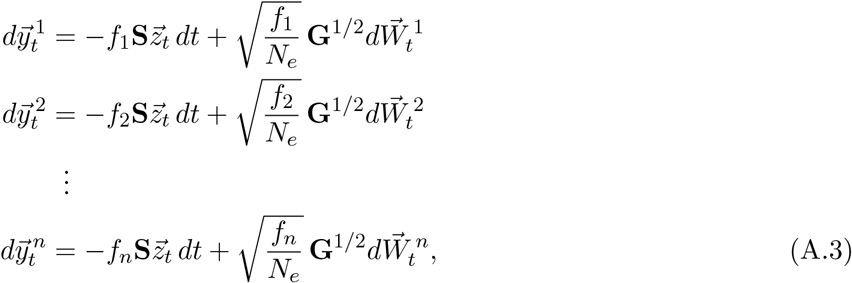

where **S** is an *m × m* matrix with elements 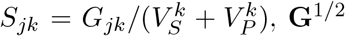 is the matrix square root and is unique and positive definite since **G** is positive definite, and the 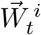 are *n* independent standard *m*-dimensional Brownian motion.

Summing the equations in (A.3), we find that the overall mean trait values obey

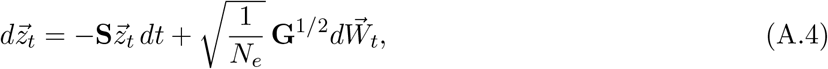

where 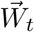 is an *m*-dimensional standard Brownian motion. 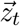 therefore behaves as a multivariate OU process (Lande 1979); its trajectory is a multivariate Gaussian process described by

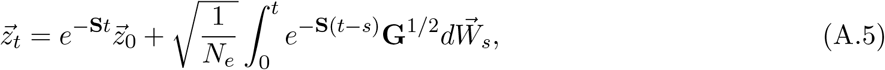

where we have used the matrix exponential. The distribution of 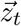 converges to 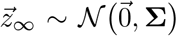, where the variance matrix **Σ** satisfies **SΣ** + **ΣS**^⊤^ = **G***/N*_*e*_.

To isolate the behavior of an individual component 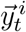, we write 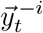 for the sum of all components other than *i*. Then, from Eq. (A.3),

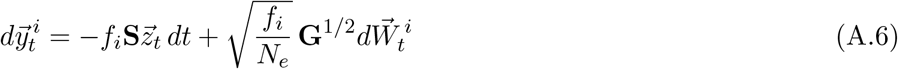

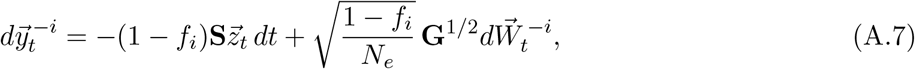

Where 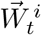 and 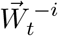 are independent standard *m*-dimensional Brownian motions. Note that, since Eqs. (A.6) and (A.7) sum to Eq. (A.4), we have

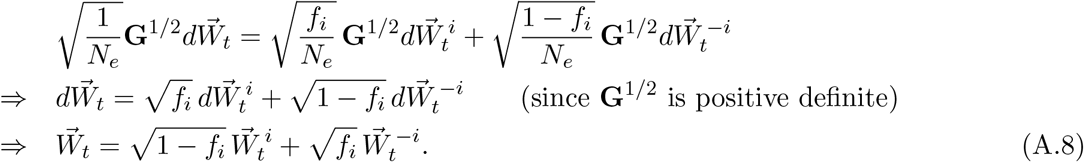

We now divide (A.6) by *f*_*i*_ and (A.7) by 1 − *f*_*i*_, subtract the latter quotient from the former, substitute in 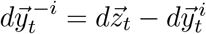, and multiply through by *f*_*i*_(1 − *f*_*i*_) to find

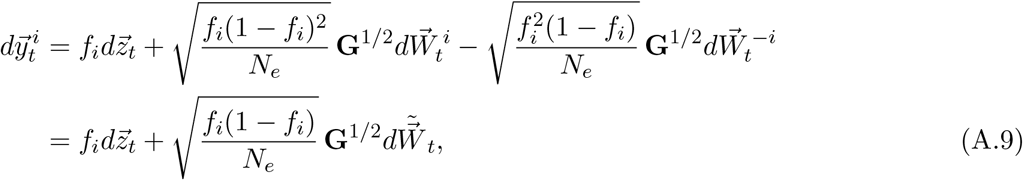

where 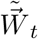 is a standard *m*-dimensional Brownian motion related to 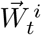 and 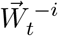 by

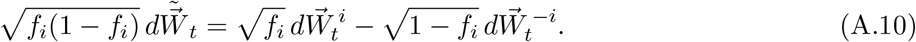

To solve for the trajectory of 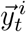, we integrate Eq. (A.9):

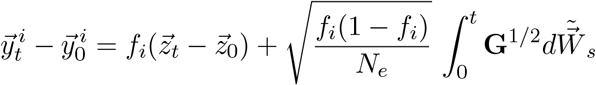

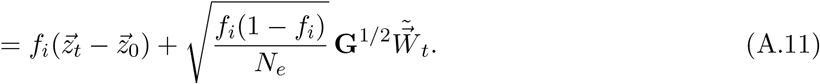

The first term in Eq. (A.11), 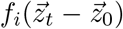, is a multivariate OU process describing the movement of the overall mean value to its optimum, and its subsequent small fluctuations around the optimum (Eq. A.5). The second term, 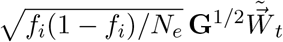, is a multivariate Brownian motion.

We now show that the two processes described by the two terms on the right-hand side of Eq. (A.11) are independent. Their stochastic components are multiples of 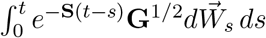 (Eq. A.5) and 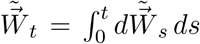 (Eq. A.11) respectively, and so we must show that these two stochastic integrals are independent. We first show that they are jointly Gaussian, and then show that they are uncorrelated, establishing their independence.

For any *m × m* matrices **A** and **B**, the linear combination

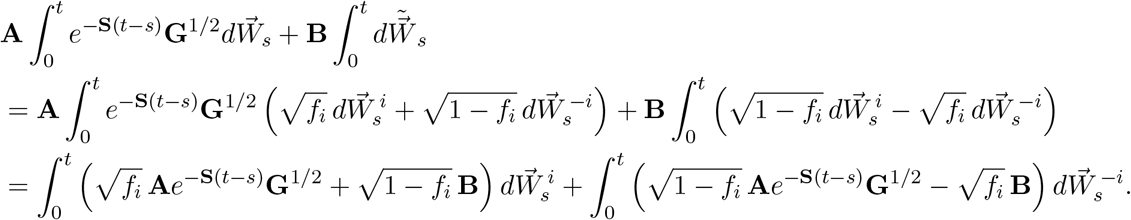

The two terms in the last line are linear functionals of independent Brownian motions 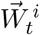 and 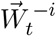 respectively, and are therefore independent Gaussian processes; their sum is therefore Gaussian. Therefore, any linear combination of the stochastic integrals in Eq. (A.11) is Gaussian, and so they are jointly Gaussian. We now show that their covariance is zero:

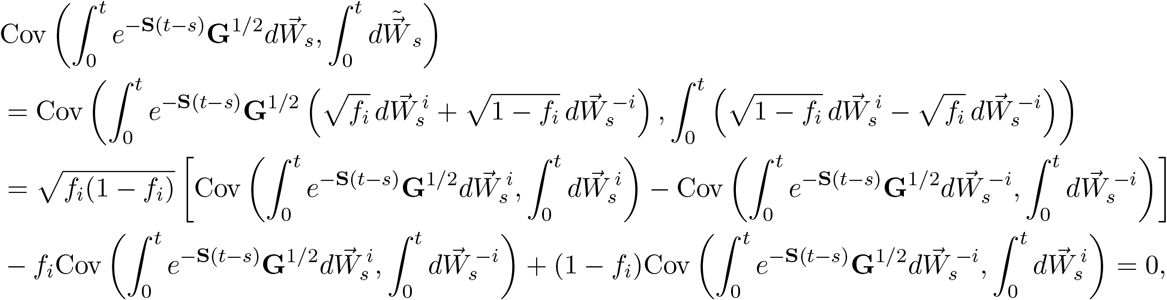

where the two covariances in the penultimate line cancel each other because of the symmetry of 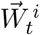 and 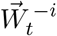, and the two covariances in the last line are each zero because of the independence of 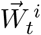 and 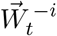.

The two terms on the right-hand side of Eq. (A.11) are thus jointly Gaussian and uncorrelated, and they are therefore independent. 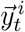 is therefore a multivariate Gaussian process with mean

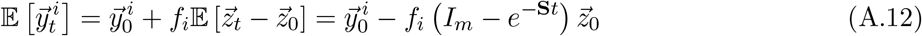

(where *I*_*m*_ is the *m × m* identity matrix) and variance

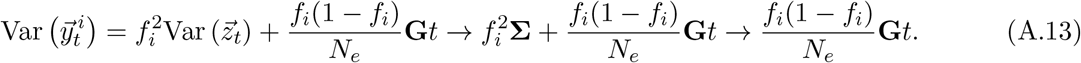

## A3 Subregion drift with recurrent migration between two populations

Let 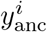 and 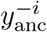 be the mean genetic values for genomic regions *i* and −*i* in the ancestral population, and let 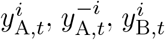, and 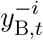 be the corresponding values in populations A and B *t* years after they split from the ancestral population. The genome-wide genetic values are 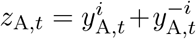 and 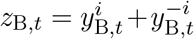. We assume for simplicity that the effective sizes of populations A and B are the same (*N*_*e*_), as are their phenotypic variances *V*_*P*_, and that their genetic variances in genomic regions *i* and −*i* are the same, with region *i* accounting for a fraction *f*_*i*_ of the total genetic variance *V*_*G*_ in each population. Each generation, a fraction *m* of population A migrates to population *B* and vice versa. In this case, the mean genetic values of the two populations in genomic regions *i* and −*i* evolve according to

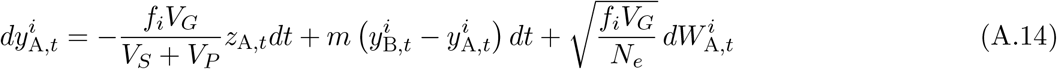

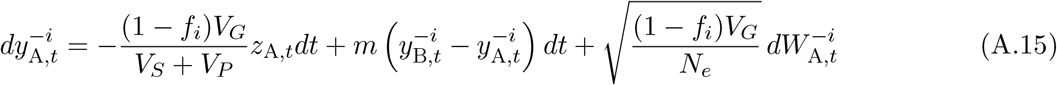

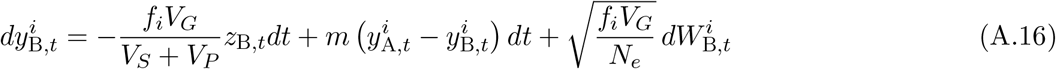

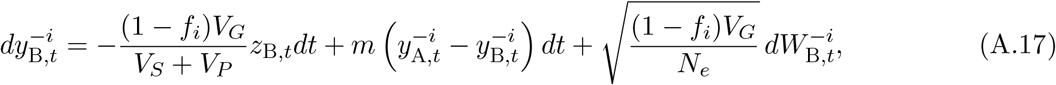

where 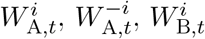, and 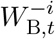 are independent standard Brownian motions. From these,

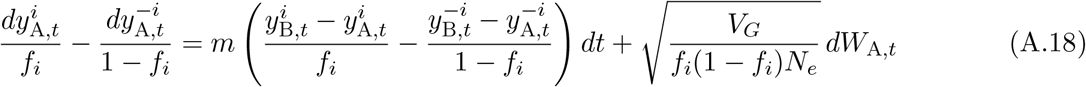

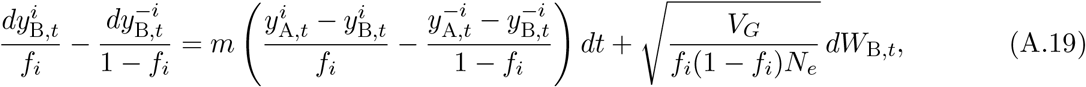

where 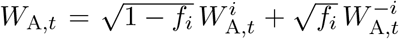 and 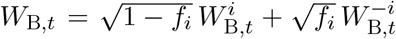 are independent standard Brownian motions. Subtracting (A.19) from (A.18),

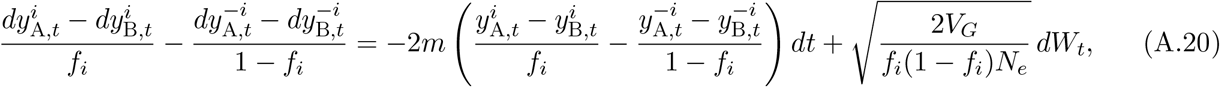

where *W*_*t*_ = (*W*_A,*t*_ − *W*_B,*t*_)*/*2 is a standard Brownian motion. But 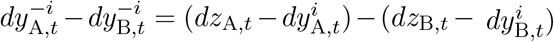 and 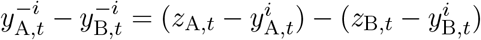, which we substitute into (A.20) to find

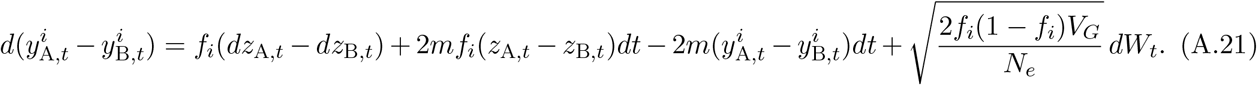

The first two terms on the right-hand side of Eq. (A.21) concern the difference between the populations in their genome-wide mean values for the trait, and can therefore be ignored in the long run. This leaves

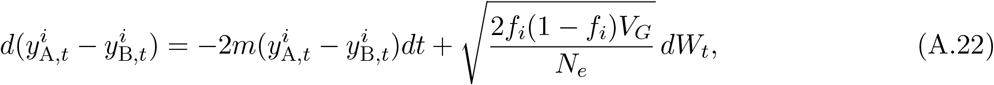

an OU process whose stationary distribution is normal with mean 0 and variance *f*_*i*_(1 − *f*_*i*_)*V*_*G*_*/*2*N*_*e*_*m*.

## A4 Full selection gradients in Application 2: Public goods production in an interspecific mutualism

There are two species involved in the mutualism. Each period, or generation, individuals from species 1 pair up with individuals from species 2. If, in a given pairing, the species-1 and species-2 partners produce amounts *γ*_1_ and *γ*_2_ of the public-good chemical respectively, then the species-1 partner has relative fitness

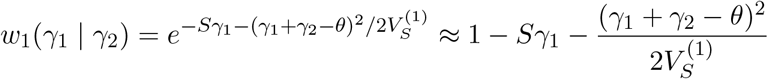

(to be compared against the fitnesses of other members of species 1), while the species-2 partner has relative fitness

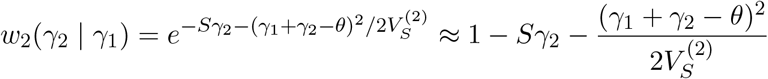

(to be compared against members of its species). In these expressions for relative fitness, the first term in the exponent is the reduction in fitness due to the cost of producing the chemical, and the second term in the exponent is the reduction in fitness due to stabilizing selection on the amount of the chemical produced by the interacting partners.

The fitness gradient associated with directional selection against chemical production is −*S*. To derive the fitness gradient associated with stabilizing selection, we assume that overall selection is weak enough that stabilizing selection’s effect on the trait mean can be considered independently of that of directional selection. Write the fitnesses due to stabilizing selection as

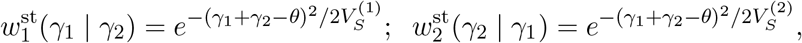

and suppose that, in a given generation, the amount of chemical produced by individuals in in each species is normally distributed, with mean *y*_1_ and variance 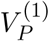 in species 1, and *y*_2_ and 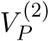 in species 2. The mean relative fitness due to stabilizing selection of a species-1 individual producing *γ*_1_ of the chemical, taken across the distribution of potential species-2 partners, is then

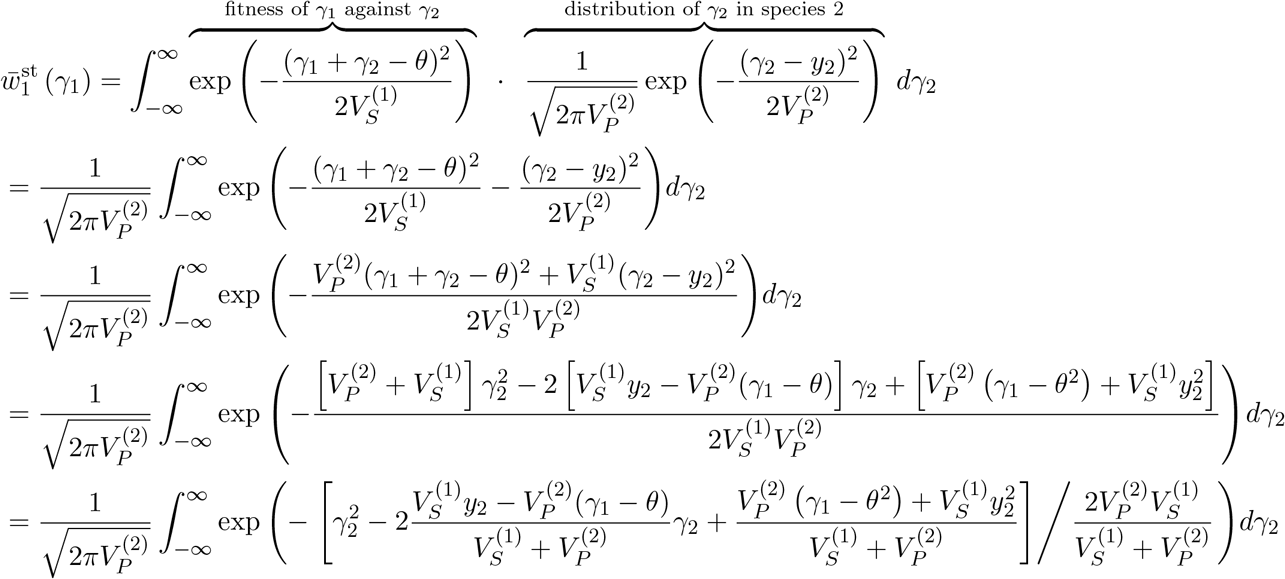

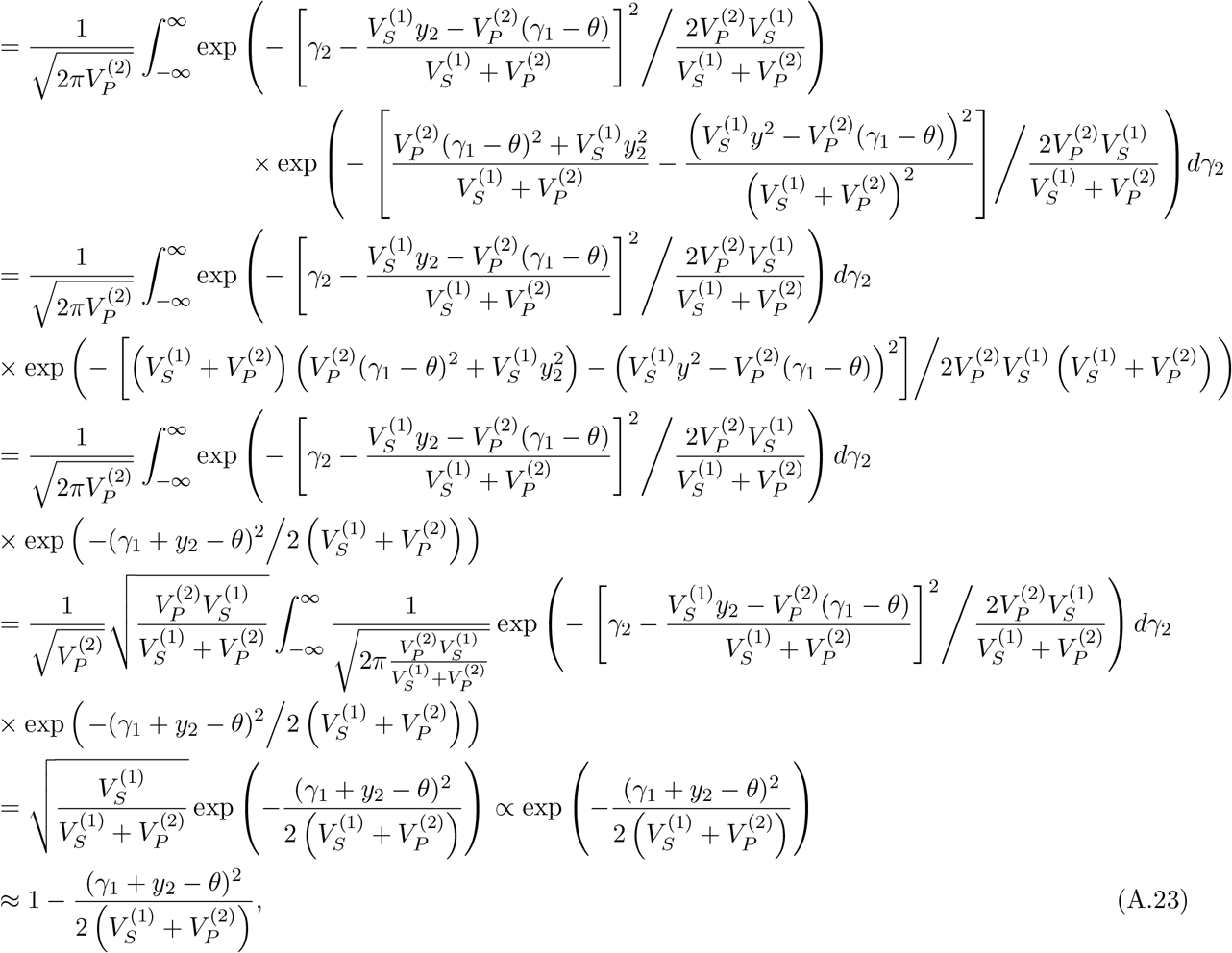

where the elimination of the integral follows from the fact that the integrand in that step is the density function of a normally distributed random variable (with mean 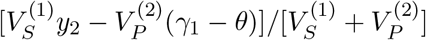 and variance 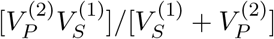).

The mean fitness in species 1 due to stabilizing selection is then

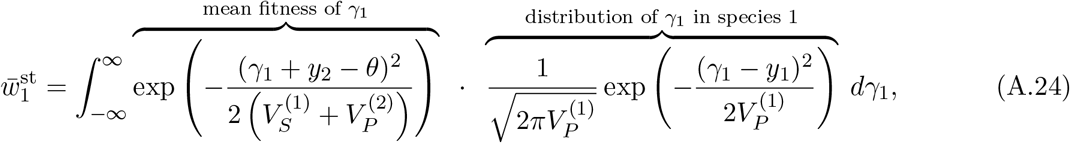

which, by techniques identical to those used in the calculation of 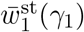 above, can be shown to be approximately

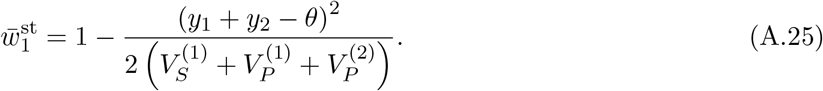

Since we have assumed the contributions of members of species 1 to be normally distributed, we can use the formulation of Lande (1976) to find the selection gradient due to stabilizing selection in species 1:

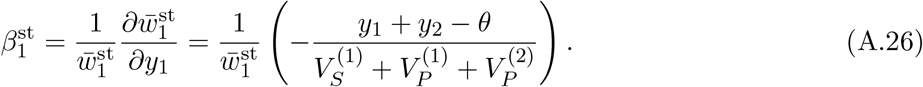

If stabilizing selection is weak 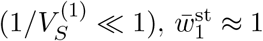, and

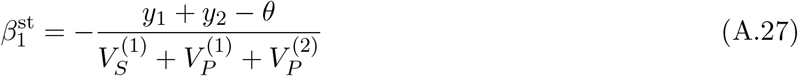

to first order in 1*/V*_*S*_. Similarly, the selection gradient due to stabilizing selection in species 2 is, to first order in 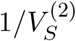,

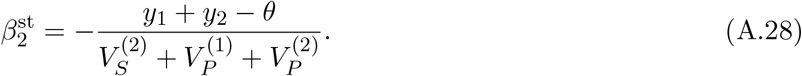

## A5 Sex chromosome drift

At locus *l*, which could be autosomal, X-linked, or Y-linked, the reference allele is at frequency *p*_*l*_ and has haploid effect 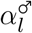 in males and 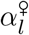 in females, relative to the alternative allele’s effects of zero. The mean haploid contribution of the locus in sex *i* is therefore 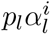, and so the change in this mean contribution across a single generation is 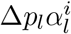.

Under stabilizing selection, the selection gradient in males is 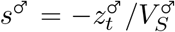 while that in females is 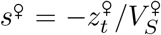, where 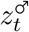 and 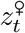 are the mean trait values in males and females in generation *t*, and 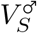 and 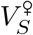 are the (inverse) strengths of selection in males and females.

### Autosomal loci

If locus *l* is autosomal, the within-generation changes in frequency at *l* in males and in females, owing to selection on the trait, are

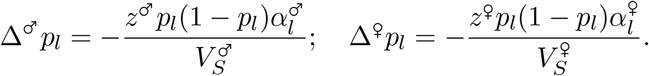

Since males and females carry an equal number of autosomes, the overall change in frequency at the locus is

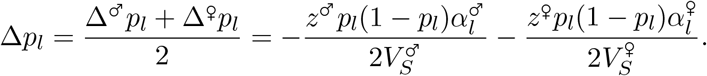

The change in the mean male genetic value at the locus is

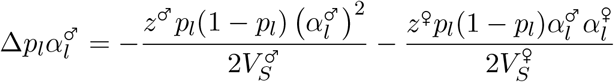

and the change in the mean female genetic value at the locus is

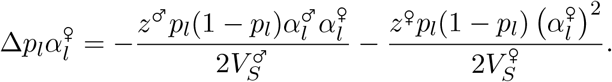

Across the set of all autosomal loci *L*_*A*_, these changes sum to

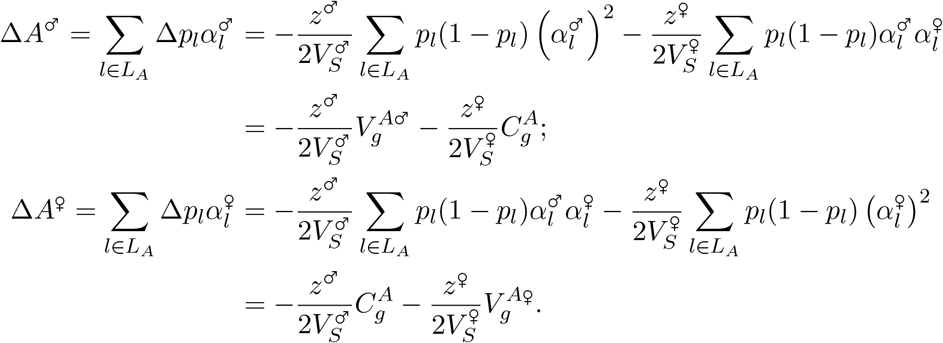

### X-linked loci

If locus *l* is X-linked, the within-generation changes in frequency at *l* in males (haploid at the locus) and in females (diploid at the locus), owing to selection on the trait, are

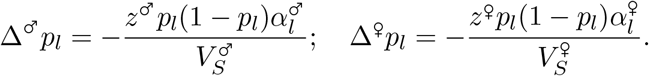

Since females carry 2*/*3 of X chromosomes, the overall change in frequency at the locus is

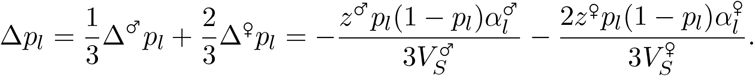

The change in the mean male genetic value at the locus is

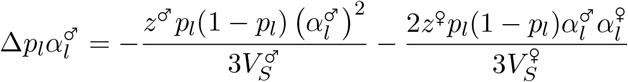

and the change in the mean female genetic value at the locus is

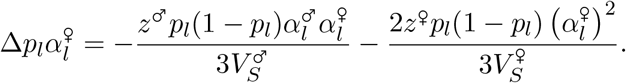

Across the set of all X-linked loci *L*_*X*_, these changes sum to

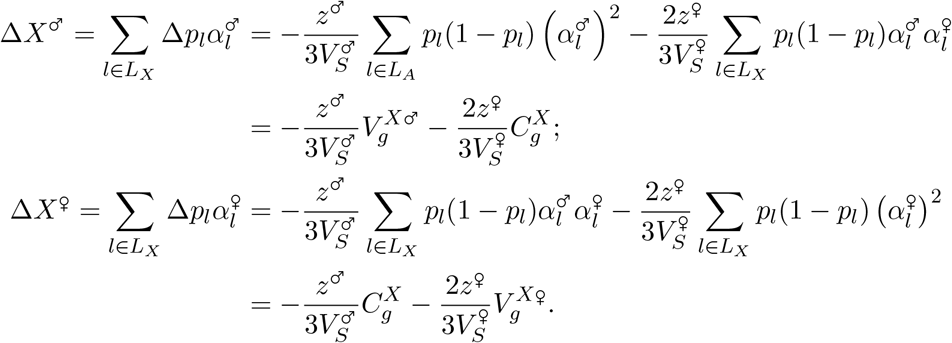

### Y-linked loci

If locus *l* is Y-linked, the within-generation change in frequency at *l* in males (haploid at the locus), owing to selection on the trait, is

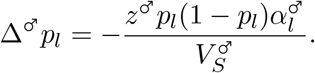

Since only males carry Y chromosomes, this is also the overall change in frequency at the locus:

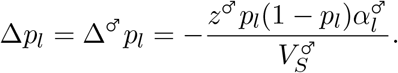

The change in the mean male genetic value at the locus is

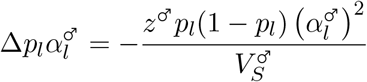

and the change in the mean female genetic value at the locus is

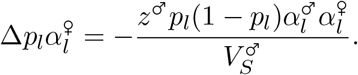

Across the set of all Y-linked loci *L*_*Y*_, these changes sum to

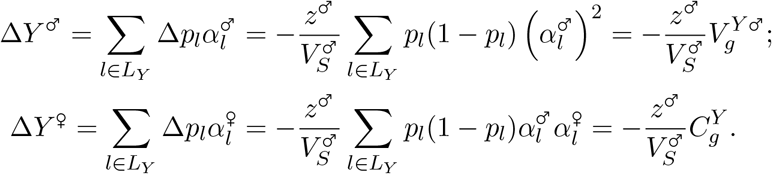

### A6 The effect of mutation on the long-term behavior of a system’s components

We return to the baseline model, with *n* genetically variable components that contribute additively to a quantitative system, the value of which is under stabilizing selection. For a given component *i*, let the set of loci with causal effects on component’s contribution to the system be *L*^*i*^. Some of these loci will be polymorphic, and thus contribute genetic variation to the value of the component, while others will be fixed, contributing no genetic variation but nonetheless affecting the mean value of the component. We assume that there are two possible alleles at each locus, one relatively trait-increasing and the other relatively trait-decreasing. The effect size of the trait-increasing allele is +*α*_*l*_, such that the expected difference between the two possible homozygotes at the locus is 2*α*_*l*_. In generation *t*, the frequency of the trait-increasing allele at locus *l* is *p*_*lt*_. If the locus is fixed for the trait-increasing allele, *p*_*lt*_ = 1; if it is fixed for the trait-decreasing allele, *p*_*lt*_ = 0; if the locus is polymorphic, 0 < *p*_*lt*_ < 1. The mean contribution of component *i* at time *t* is then

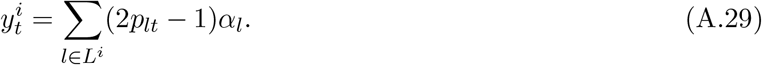

We now suppose that the two alleles at each locus can mutate to one another. We assume that the rate of mutation is *μ* per replication, symmetric between the two alleles at each locus and constant across loci. At locus *l*, the expected change in frequency of the trait-increasing allele due to mutation is then Δ^mut^*p*_*lt*_ = *p*_*lt*_(1 − *μ*) + (1 − *p*_*lt*_)*μ* = −(2*p*_*lt*_ − 1)*μ*. The expected change in the mean genetic value of the component due to mutation in generation *t* is therefore

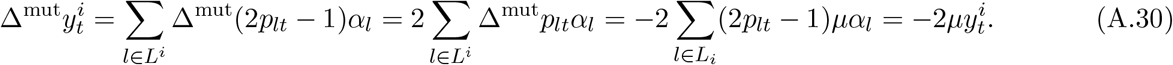

That is, components that have attained high mean values owing to the rise in frequency, and fixation, of trait-increasing alleles will tend to have their values decreased by mutation, because most mutations affecting these components will change (high-frequency) trait-increasing alleles to trait-decreasing alleles. Mutation will thus tend to check the spread of components’ contributions to the system as they get far from their initial values.

Incorporating Eq. (A.30) into Main Text Eqs. (6) and (7), we find that under mutation, selection, and drift, the mean value of the *i*-th component and the sum of the mean values of all other components obey

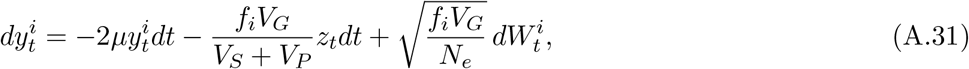

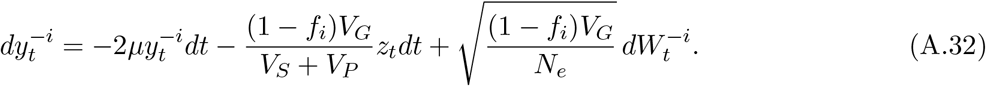

The mean genetic value of the overall system obeys

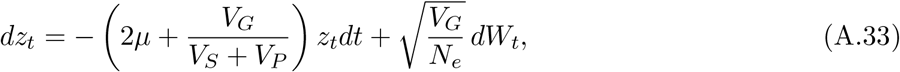

which can be obtained by summing Eqs. (A.31) and (A.32). *z*_*t*_ is therefore an OU process. Note that the per-locus mutation rate *μ* will usually be much smaller than *V*_*G*_*/*(*V*_*S*_ + *V*_*P*_)—for instance, in mutation-selection-drift equilibrium under stabilizing selection, *V*_*G*_*/V*_*S*_ ≈ 4*Lμ*, where *L* is the number of loci at which new mutations would affect the trait (e.g., Turelli and Barton 1990). In Eq. (A.33), *μ* ≪ *V*_*G*_*/*(*V*_*S*_ + *V*_*P*_) implies that selection is a stronger force pulling the mean system value towards its optimum than selection is. The trajectory of *z*_*t*_ under Eq. (A.33) is

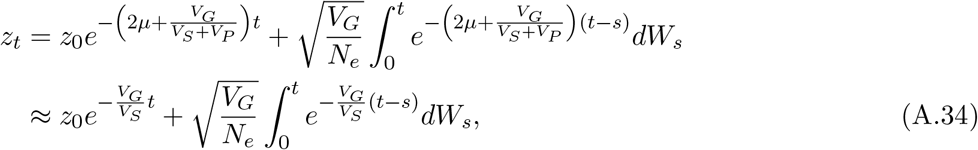

where the approximation follows from *μ* ≪ *V*_*G*_*/*(*V*_*S*_ + *V*_*P*_) and the fact that, under reasonable strengths of stabilizing selection, *V*_*P*_ ≪ *V*_*S*_. *z*_*t*_ is therefore a Gaussian process with mean and variance

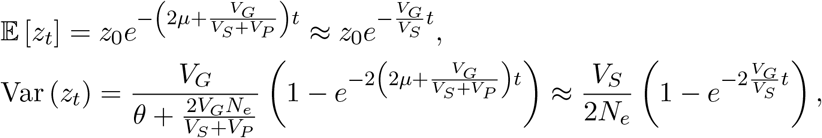

where *θ* = 4*N*_*e*_*μ* is the population-scaled per-locus mutation rate. The stationary distribution of *z*_*t*_ is normal with mean zero and variance

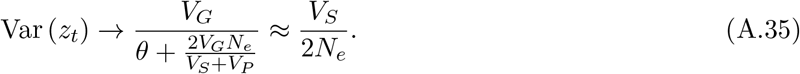

To isolate the behavior of an individual module 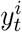, we proceed as before. We first divide Eqs. (A.31) and (A.32) by *f*_*i*_ and 1 − *f*_*i*_ respectively:

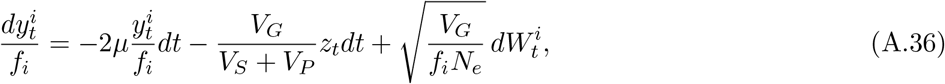

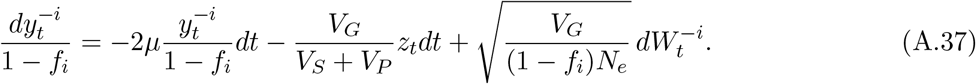

We then subtract (A.37) from (A.36):

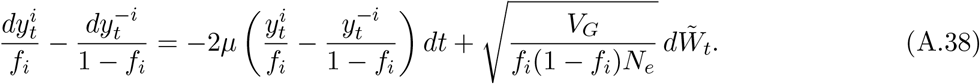

Finally, we substitute 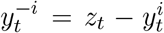 (and 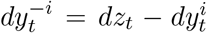) into Eq. (A.38) and multiply through by *f*_*i*_(1 − *f*_*i*_) to obtain

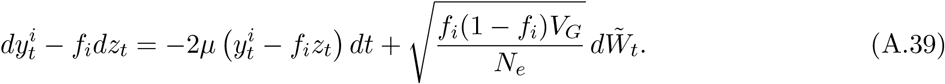

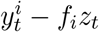 is therefore an OU process, the trajectory of which can be written

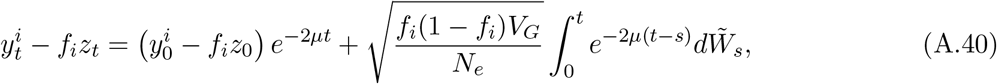

which is a normal random variable with mean 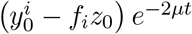 and variance 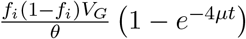. The stationary distribution of 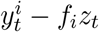 is therefore 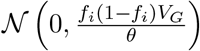.

From Eqs. (A.34) and (A.40), the trajectory of the mean value of component *i* is

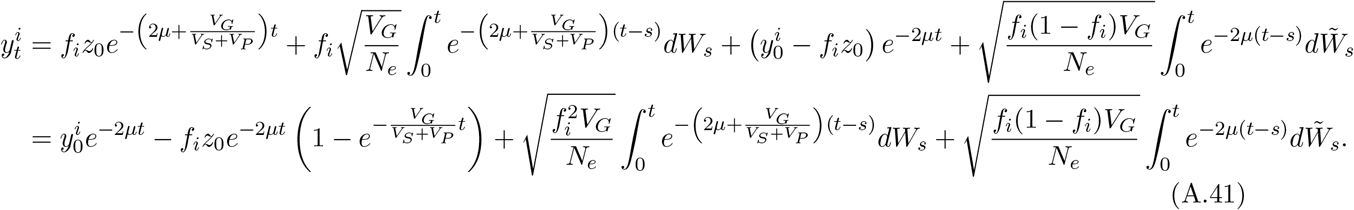

The two stochastic integrals in Eq. (A.41) are Gaussian and independent (following the same logic as in Appendix A1), and so 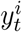 is a Gaussian process with mean

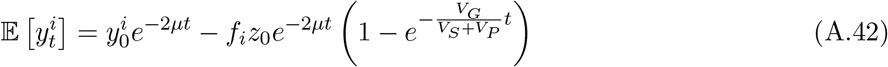

and variance

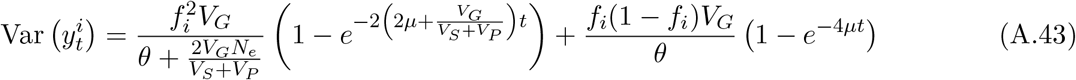

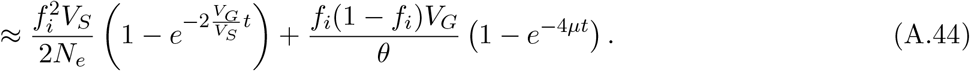

This converges to a stationary distribution which is normal with mean 0 and variance

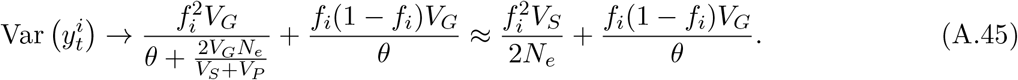

Under stabilizing selection, in mutation–selection–drift balance in the large-population limit, *V*_*G*_ ≈ 4*V*_*S*_*Lμ*, where *L* is the number of loci at which mutations affect any of the components in the system (e.g., Turelli and Barton 1990). Substituting this value into Eq. (A.45), we find

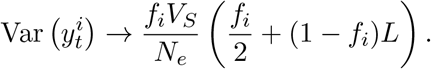

The second term in the brackets, (1 − *f*_*i*_)*L*, is much larger than the first, *f*_*i*_*/*2, as long as component *i* contributes substantially more than 1*/L* of the system’s genetic variance—i.e., more than a single locus’s worth. Therefore, we can take the long-term variance of component *i*’s mean contribution under the system’s mutation–selection–drift balance to be

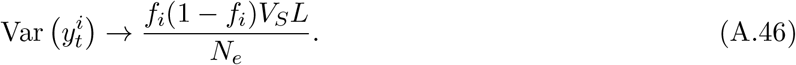

In contrast, from Eq. (A.35), the long-term variance of the overall system’s mean value is

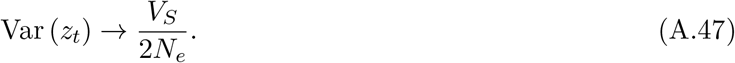

The ratio of component *i*’s long-term variance with the overall system’s is therefore

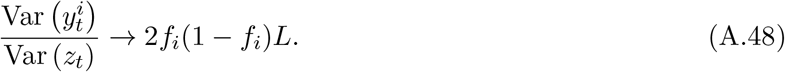

Therefore, unless component *i* contributes almost none (∼ 1*/L*) or almost all (∼ 1 − 1*/L*) of the genetic variance of the system, the variance of the stationary distribution of its mean will be much larger than that of the system’s. That is, in the long-term, the mean contribution of component *i* to the system will drift around in a very broad range of values, despite the system’s mean value being constrained to a very narrow range around the optimum.

## Footnotes

1 To see that the 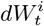 terms in Eq. (5) combine to give *dW*_*t*_ in Eq. (1), note that 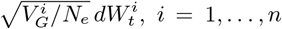, are independent normal random variables with zero mean and variance 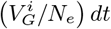; their sum is therefore a normal random variable with zero mean and variance 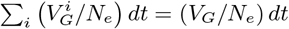.

2 For highly polygenic traits, the fraction of the genetic variance contributed by a given genomic region is often approximately equal to the region’s fraction of total genome length (Yang et al. 2011).

3 The assumption that the optima are the same for males and females is not as restrictive as it might appear since, with different optima and imperfectly correlated allelic effects, males and females would rapidly move to their respective optima, after which stabilizing selection would resume with genetic dynamics that are the same as if the optima were identical in the first place (Muralidhar and Coop 2024).

